# Transposable Element Diversification and the Evolution of Peltigerales Lichen Symbionts

**DOI:** 10.64898/2026.02.14.705750

**Authors:** Ellen S. Cameron, Benjamin J.M. Tremblay, Rebecca Yahr, Mark L. Blaxter, Robert D. Finn

## Abstract

Lichens are composite organisms formed through the symbiotic association between fungi, algae and/or bacteria. Multiple independent origins of the lichenized lifestyle have been reported in both fungal and algal lineages, but the molecular mechanisms and evolution underpinning these symbiotic relationships remain largely unknown. In this study, we performed long-read metagenomic sequencing on 11 Peltigerales lichen species to characterize the genomic content of lichen symbionts via metagenome assembled genomes (MAGs). Peltigerales genomes generated in this work represent the largest Lecanoromycetes genome sequenced to date, driven by high transposable element content. Transposable elements (TEs) are known to drive genome evolution in other symbioses but have been underexplored in lichen symbionts due technological limitations. Transcriptomics revealed that many genes associated with adaptations to the lichenized lifestyle are associated with TEs suggesting that they may play a key role in the evolution of lichenization.

## 1. Introduction

Lichens have traditionally been viewed as the symbiotic association between a fungal partner and photosynthetic algal partner^1^. It is estimated that there are over 20,000 species of lichenized fungi^2^, with most forming symbiotic associations with chlorophyte algae. However, some lichens are formed through the symbiosis between a fungal partner and a cyanobacterium (‘cyanolichens’), and in other cases both chlorophyte and cyanobacterial partners are involved (‘tripartite cyanolichens’)^3,4^. Multiple independent origins of lichenization are inferred in diverse fungal lineages^5,6^, and in cyanobacterial^7^ and chlorophyte photobionts^8^. More recently, the diversity of what constitutes a lichen has expanded with metagenomic surveys revealing complex multispecies assemblages beyond primary fungal and algal partners, consisting of diverse bacterial communities^9,10^, additional fungi^11,12^, and viruses^13^. However, fundamental questions of the function and evolution for the two primary symbionts of the lichen, fungi and algae, remain unanswered^1,14^. Long-read metagenomic sequencing provides new avenues to explore these questions further through the assembly of high-quality metagenomic assembled genomes (MAGs), which can be used to explore the genomic characteristics of symbionts suggested to drive the evolution in other fungal interactions.

Transposable elements (TEs), also known as transposons or mobile elements, can move within a genome via transposition and often represent the majority of repetitive sequences found in eukaryotic genomes^15,16^. In fungi, TEs may contribute to key roles in adaptation to symbiotic or pathogenic lifestyles^17,18^ with TEs previously being found to be involved with genes associated with symbiosis in ectomycorrhizal fungi^19^ or virulence factors in pathogenic fungi^18,20^. Proliferation of TEs can result in expansions in genome size and the generation of new mutations, but mechanisms exist to repress TE activity including epigenetic silencing and loss through degradation^21^. In rare cases, TE insertions may be beneficial to their hosts, leading to domestication and retention in the host genome^21^ or contribute to genetic variability driving the coevolution between symbionts^22^. The proportion of eukaryotic genomes made up of TEs ranges from only 3% in yeast to upwards of 80% in some plants^17^. However, in some fungi with host-associated lifestyles, TEs contribute to large proportions of the total genome size as observed in arbuscular mycorrhizae (47%)^18^ or the fungal pathogen, *Austropuccinia psidii* (>90%) which has the largest fungal genome sequenced to date^19^. Despite TE content being identified as a potential driver of coevolution in plant-associated fungi with diverse lifestyles ranging from pathogenic to mutualism, few papers have addressed the role of TEs in other fungal symbioses including lichens^23,24^. The ability to explore the role of TEs rely on high-quality, highly contiguous genomic assemblies which may not be consistently achieved with short-read shotgun metagenomics^9^.

In this study we used long-read metagenomics and short-read transcriptomics to characterize all the organismal constituents and their associated genomic content, including TE content, for 11 different species of cyanolichens. We generated highly contiguous metagenomic assembled genomes (MAGs) for primary fungal symbionts and photobionts of cyanolichens, including the first genome assemblies for ten of the lichenized fungal species. Using these MAGs, we demonstrated that TE content may contribute to the evolution in Peltigerales fungi by (1) comprising a large portion of the genomic content and (2) being associated with highly expressed genes associated with stress response and cellular communication.

## 2. Methods

### 2.1 Sample collection, nucleic acid extraction & sequencing

Lichen thalli of 11 cyanolichen species (Table S1) were collected with permission from Glen Creran Woods Site of Special Scientific Interest (Argyll, Scotland) and air dried for preservation for 72 hours. Associated bryophyte material was removed from lichen thallus prior to long-term storage in a −80°C freezer.

Thallus tissue for samples used in analysis were prepared for nucleic acid extraction using a CryoPrep for flash frozen tissue pulverisation. The resulting pulverised tissue powder was used for DNA extraction using the protocol proposed by Park et al. (2014)^25^, which utilises a potassium chloride extraction buffer with chloroform (Table S2). DNA extracts were assessed for quantity and quality using a NanoDrop spectrophotometer and Qubit fluorometric assay. DNA extracts were sheared using a MegaRuptor at a speed of 130, and SPRI cleaned up (0.88 v/v concentration) with the addition of RNase to degrade any remaining RNA material. Sheared DNA extracts were assessed for suitable fragment length using the Agilent FemtoPulse. Final DNA extracts were submitted for PacBio HiFi sequencing on the Revio platform at Scientific Operations at the Wellcome Sanger Institute.

Thallus tissue from two individuals or two parts of a single individual of each species used for genomic sequencing were pulverized for RNA extraction (Table S3). RNA extractions were performed following the manufacturer’s protocol for TRIzol including a second chloroform wash step prior to isopropanol precipitation to remove phenol contamination. RNA extracts were treated with DNase using the Turbo DNase kit to remove remaining DNA. Resulting RNA extracts were assessed for quantity and quality using an Agilent RNA Screen Tape Station analysis and were submitted for NEB NextERA RNAseq Illumina sequencing on the NovaSeqX platform at Scientific Operations at the Wellcome Sanger Institute.

### 2.2 Generation of metagenomic assembled genomes (MAGs)

PacBio HiFi reads were assembled into longer contiguous sequences using *metaMDBG* (v. 0.3; Table S4)^26^. Reads were mapped back to resulting metagenomic assemblies using *minimap2* (v. 2.23)^27^ to determine assembly coverage. Contigs were binned into representative genomes using *CONCOCT* (v. 1.1.0)^28^ and *metaBAT2* (v. 2.12.1)^29^.

Eukaryotic bins were identified from the set of bins generated with *metaBAT2* with *EukCC* (v. 2.1.2)^30^. Eukaryotic bins were assessed for completeness using *EukCC* and *BUSCO* (v. 5.7.0; Table S5; Table S6)^31^, and merged bins were generated using *EukCC* where applicable.

To explore the bacterial portion of the lichen metagenome, bins from *CONCOCT* and *metaBAT2* were refined using the *metaWrap* (v. 1.3)^32^ bin refinement module. Refined bin sets were dereplicated using *dRep* (v. 3.2.2; Table S7)^33^ and taxonomically classified using *GTDBtk* (v. 2.3.0)^34^. Quality scores (QS; completeness - [5 * contamination]) were calculated for bacterial bins using metrics determined by *CheckM* (v. 1.1.3; Table S8)^35^. Bacterial bins with a QS > 50 were retained for gene annotation with *prokka* (v.1.14.6)^36^. Predicted genes were used for functional annotations with *InterproScan* (v. 5.60-92.0)^37^. For phylogenetic tree reconstruction, ModelFinder^38^ was used to identify the best-fit model according to the Bayesian Information Criterion (BIC). Subsequently, a phylogenetic tree for bacterial MAGs was generated using alignments from *GTDBtk* with *iqtree* (v. 2.2.0)^39^ using model LG+F+I+G4 with 1000 bootstraps and visualized with *ggtree* (v. 3.12.0)^40^

Genomic composition of metagenomes was assessed by mapping reads with *minimap2* to assemblies and calculating a weighted mean depth as a function of the length of the MAG (Table S9) and contig to the number of reads mapped. MAGs were identified as being ‘present’ in a metagenome if >30% of the MAG was covered by reads (Table S10; Table S11).

### 2.3 Exploration of cyanobacterial symbiont genomes

Synteny analysis of cyanobacterial MAGs was performed using *jcvi*^41^ to identify genomic arrangements between different detected *Nostoc* species. KEGG orthologs identified with KofamScan (v. 1.3.0)^42^ annotations corresponding to nitrogen fixation pathways were identified in cyanobacterial MAGs to explore the occurrence of iron-molybdenum dependent and vanadium dependent nitrogen fixation pathways in cyanobacterial symbionts. Cyanobacterial MAGs were placed into a phylogenetic tree with other reference genomes (Table S12) to explore habitat associations. Cyanobacterial MAGs and reference Nostocaceae and Leptolyngbyaceae genomes were dereplicated using *dRep* (Table S13). Alignments were generated for the dereplicated set of cyanobacterial genomes, including both references and MAGs, using GTDBtk. ModelFinder identified the best-fit model according to the BIC and a phylogenetic tree created with *iqtree* using model LG+F+R8 with 1000 bootstraps and visualized with *iTOL* (v. 7)^43^.

### 2.4 Annotation and characterization of eukaryotic symbionts

Genome assemblies for fungi (Lecanoromycetes; Table S14) and green algae (Chlorophyta; Table S15) were downloaded from NCBI in June 2024 and retained for analysis if the N50 was greater than 1Mb to ensure the retention of high-quality genome assemblies. Phylogenetic trees were generated for fungal and algal lineages using BUSCO phylogenomics (https://github.com/jamiemcg/BUSCO_phylogenomics) in which BUSCO groups shared amongst 90% of genomes are concatenated and aligned for placement. ModelFinder was used to identify the best model according to BIC. A phylogenetic tree was constructed for the Lecanoromycetes assemblies using the JTT+F+I+G4 model with 1000 bootstraps. For the Chlorophyta assemblies, the LG+F+I+G4 model was used for phylogenetic tree construction with 1000 bootstraps in *iqtree*. Telomeres were identified in eukaryotic MAGs using a script from Hiltunen *et al.*, with a query search of TAA[C]+ for fungal genomes and CCCATTT for algal genomes as described in Tagirdzhanova *et al.*^44,45^ GC content was calculated per contig with a sliding window size of 10,000 using *universalmotif* (v. 1.22.3)^46^.

To further explore the taxonomic identity of the chlorophyte photobiont, 18S rRNA gene sequences were identified in the recovered chlorophyte photobiont MAG with *barrnap* (v. 0.9)^47^ and placed phylogenetically with other 18S rRNA gene sequences from other lichenized and non-lichenized chlorophyte algae (Table S16) using the TNe+I+G4 model as identified by ModelFinder in *iqtree*.

Putative organellar genomes for the chlorophyte photobiont species were identified using *tiara* (v. 1.0.3)^48^ classifications and confirmed based on:(i) size;(ii) circularity; and (iii) gene annotation. Chloroplast genomes for chlorophyte photobionts were assessed for completeness using quality scoring algorithms present in *plastiC* (v.0.1.2)^49^. Mitochondrial and chloroplast genomes were annotated and visualized with *OG-DRAW* (v. 1.3.1)^50^.

Initial protein predictions were performed on eukaryotic MAGs using *metaEuk* (v. 7-bba0d80)^51^ with a reference database corresponding to respective lineages of Trebouxiophyceae for chlorophyte photobionts and Lecanoromycetes for lichenized fungi. These protein predictions were used alongside MAGs to characterize splicing characteristics and were used to generate an index with *STAR* (v. 2.7.11b)^52^ for each individual species. Raw RNAseq reads were trimmed using *fastp* (v. 0.23.4)^53^ and mapped to MAGs using *STAR. stringtie* (v. 2.2.3)^54^ was then used to predict transcripts based on this mapping, using BAM files produced by *STAR* split into forward and reverse strands using *samtools* (v. 1.21)^55^ to improve the annotation of overlapping genes. Final transcript sequences were generated with *gffread* (v. 0.12.7)^56^ and were used for coding sequence (CDS) generation with *Transdecoder* (v. 5.7.1)^57^ including homology searches conducted with *hmmer* (v. 3.1b2) and *diamond* (v. 2.0.14)^58^. Gene expression was quantified using *salmon* (v. 1.10.3)^59^.

For samples with more than one eukaryotic MAG present (including tripartite cyanolichens, and those with additional fungi), eukaryotic bins and GTF files for each detected species were merged to generate a combined species index with *STAR* for mapping, and subsequent steps were followed as highlighted above.

The gene annotations as generated above were functionally annotated with *InterProScan*.

### 2.5 Characterizing transposable element content

Repeat content of eukaryotic genomes was characterized using *EarlGrey* (v. 5.1)^60^. Transposable elements less than 100 bp in size were discarded from downstream analyses. Overall repeat content (%) of the genome was determined based on coverage and length of identified transposable elements. The co-occurence of transposable elements and genes were visualized using *Circlize*^61^. Kimura distances, as determined by *EarlGrey*, were used to characterize the TE activity to identify TE acquisition events in the repeat landscape based on similarity to consensus sequences.

Transposable element position was used in reference to gene positions to determine genes located within or in close proximity to transposable elements. Distances of genes to transposable elements were classified into categorical bins as follows: 0bp (e.g., gene inside TE, fragments of TEs inside genes), 1-1,000bp, 1,001-5,000bp, 5,001-10,000bp, 10,001-20,000bp, >20,000 bp. To identify gene-rich or TE-rich regions, contigs were split into 25 kb windows, sliding every 5 kb. Windows were assigned as gene-rich if they contained over 60% genes, and TE-rich if they contained over 60% TEs. Overlapping windows with identical assignments were merged together.

Gene expression was explored to identify correlation of highly expressed genes, proximity to transposable elements, and functional relevance. Three approaches were used to examine the relationship between gene expression and TE proximity. First, the average log of the normalized values of transcripts per million (TPM) obtained from *salmon* for each species with the distance between each gene with their nearest TE were plotted as scatter plots. Second, the average log TPM values per species were grouped into the previously determined gene-rich, TE-rich, or mixed categories and plotted as boxplots. Third, the distribution of expressed (average log TPM value greater or equal to 1) and non-expressed (average log TPM value less than 1) gene counts per species per categorical TE distance bins were plotted as stacked bar plots.

Functional enrichment of genes grouped by categorical TE distance bins was performed using the *topGO* (v. 2.56.0)^62^ using Gene Ontology (GO) Biological Processes (BP) annotations determined by *InterProScan* to calculate enrichment within each bin. Selected enriched GO terms (p-value < 0.05) with biological relevance were used for visualization. All enriched GO terms are available in supplementary data (Table S20/S21).

The *regioneR* (v. 1.36.0)^63^ package was used to determine whether genes are present closer or further away from TEs in the lichenized fungal genomes than expected by chance by shuffling the locations of the genes in the genomes 1,000 times and calculating the average gene to TE distance for each iteration, leading to a final P-value calculation. The same procedure was carried out to determine whether specific GO:BP annotated genes were closer or further away, only instead shuffling the annotations between existing genes with annotated GO terms. Terms closer or further away than the average were semantically enriched and clustered using the *simplifyEnrichment* (v. 1.14.1)^64^ package. The *regioneR* package was also used to test whether specific TE families are closer or further away from genes than expected by average, shuffling the family label of TEs (excluding TEs with no known family).

Candidate TE fusions were identified using TE-specific Pfam domains previously identified in fungi^65^. Transcripts were characterized as TE-derived if all identified Pfam domains within the protein sequence were TE-specific, and TE fusions if the Pfam domains were a mix of TE-specific and non-TE-specific domains.

## 3. Results

### 3.1 Cyanolichens host diverse bacterial and eukaryotic microbial communities

PacBio HiFi metagenomic sequencing was conducted for 11 cyanolichen species (Figure 1A-F) leading to the generation of high-quality metagenomic assemblies (Table S4). Metagenomic assembled genomes (MAGs) for primary fungal, cyanobacterial and chlorophyte symbionts, as well as additional fungi and bacteria were found in the lichen thallus (Table S10/S11). Following genome redundancy removal, a total of 103 bacterial, 15 fungal and 1 chlorophyte MAGs were identified (Figure 1G).

**Figure 1:**
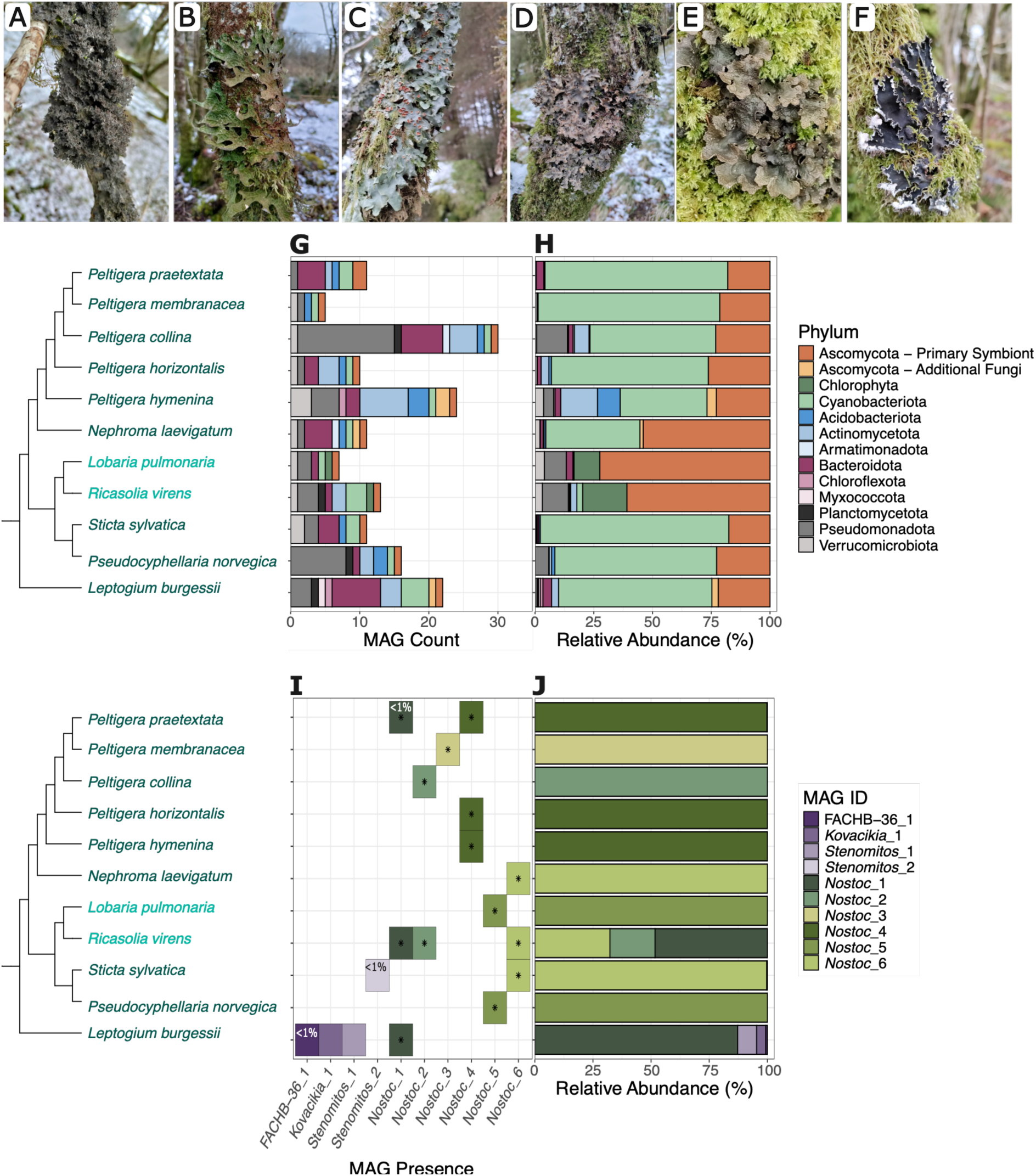
Organismal composition of 11 different species of cyanolichens determined using long-read metagenomic approaches. (A-F) Thallus of lichen species which include cyanobacterial photobionts sequenced in this study including Leptogium burgessii (A), Lobaria pulmonaria (B), Ricasolia virens (C), Pseudocyphellaria norvegica (D), Sticta sylvatica (E), and Peltigera collina (F). (G) Absolute count of detected metagenomic assembled genomes (MAGs) and (H) relative contribution of different taxa to cyanolichen metagenomic data generated. Cyanolichen species shown in turquoise (Lobaria pulmonaria and Ricasolia virens) are tripartite cyanolichens including both chlorophyte and cyanobacterial photobionts. (I) Detection of cyanobacterial MAGs belonging to Nostocaceae (green) and Leptolyngbyaceae (purple) across cyanolichen species. Cells marked with a ‘*’ indicate that nitrogen fixation pathways were detected in the genome. Genomes accounting for <1% composition of the cyanobacterial component of lichen metagenomes are indicated. (J) Relative composition of cyanobacterial components in the sequenced lichen metagenomes.

Ten bacterial phyla were represented across the 103 bacterial MAGs: Acidobacteriota, Actinomycetota, Armatimonadota, Bacteroidota, Chloroflexota, Cyanobacteriota, Myxococcota, Planctomycetota, Pseudomonadota and Verrucomicrobiota. Representatives from bacterial families commonly found in lichens including Acidobacteriaceae, Beijerinckiaceae, Acetobacteraceae and Sphingomonadaceae were detected (Figure S1). The diversity and frequency of bacterial MAGs, excluding the cyanobacterial photobiont, between samples ranged from 31 in Peltigera collina to just 3 in Peltigera membranacea. 77 MAGs were only detected in a single sample (Figure S2; Table S17). Bacteria, excluding the Cyanobacteriota, were cumulatively present in lower abundances in each lichen metagenome in comparison to the primary mycobiont and photobiont (including both chlorophyte and cyanobacteria) MAGs (Figure 1H).

Sequencing depths normalized as a function of genome/bin size were used to explore the taxonomic composition of the cyanolichen metagenome. The lichenized fungi (Ascomycota) were major components of the metagenome. Cyanobacteriota, representative of the cyanobacterial photobiont in the lichen symbioses, were frequently found to be dominant representatives within the lichen metagenome accounting for up to 80.1% by copy number (see Methods) in Sticta sylvatica, and on average comprising 51.6% of metagenomic assemblies (Figure 1H). The relative abundance of cyanobacterial photobionts in the two tripartite cyanolichens, Lobaria pulmonaria and Ricasolia virens, was significantly less (0.77% and 2.5% respectively) and chlorophyte photobionts were more abundant (10.1% and 18.1%).

### 3.2 Alternative vanadium-dependent nitrogen-fixation pathways observed in Nostoc symbionts

Of the 103 bacterial MAGs identified across the 11 species of cyanolichens, 10 were identified as cyanobacteria. These 10 cyanobacterial MAGs included six genomes identified as distinct *Nostoc* species, and 4 genomes from the family Leptolyngbyaceae. *Nostoc* species are common cyanobacterial photobionts in lichens, but Leptolyngbyaceae have not previously been reported as lichen symbionts (Figure 1I). Further exploration of the Cyanobacteriota composition revealed that Leptolyngbyaceae MAGs were present in extremely low abundance when detected (maximum 3.3% in *Sticta sylvatica*; <1% in many other instances; Figure 1J) suggesting that these cyanobacteria are unlikely to be primary photobionts. The *Nostoc* strains identified in this study showed similarity to other previously identified *Nostoc* symbionts, while Leptolyngbyaceae taxa showed similarity to references originating in aquatic sources and not from symbiotic lineages (Figure 2A).

**Figure 2:**
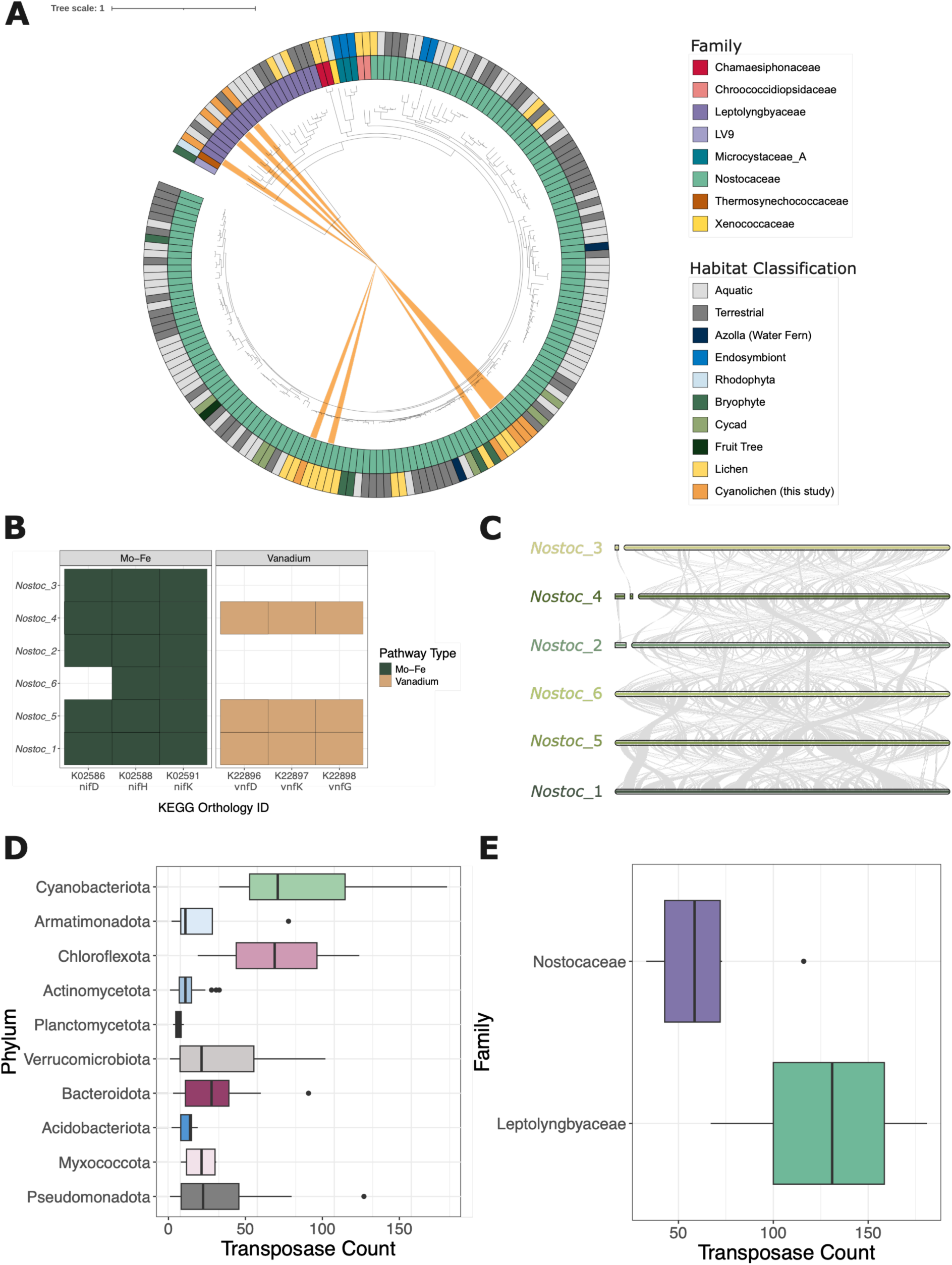
Diversity, distribution and genomic exploration of cyanobacterial metagenomic assembled genomes (MAGs). (A) Phylogenomic tree of dereplicated Nostocaceae and Leptolyngbyaceae genomes to identify conservation of symbiont types. Orange shaded branches of the tree indicate cyanobacterial MAGs generated in this study. (B) Detection of KEGG orthologs in molybdenum-iron (Mo-Fe) and vanadium dependent nitrogen fixation pathways in *Nostoc* MAGs. *vnfH* was not detected in any *Nostoc* MAG and is not shown. (C) Synteny plots of *Nostoc* genomes to identify areas of genomic rearrangements. (D) Frequency count of transposase genes in bacterial MAGs. (E) Frequency count of transposase genes in cyanobacterial MAGs only.

We found at least one nitrogen-fixing cyanobacterial symbiont per lichen thallus. However, in some cases, such as *Ricasolia virens*, multiple different species of nitrogen-fixing *Nostoc* were identified from a single thallus sample (Figure 1I). While some cyanolichens were found to host multiple species of *Nostoc*, one species was always found to be dominant in all cyanolichens assessed. Nitrogen-fixation pathways in these *Nostoc* symbionts included the molybdenum-iron dependent pathway in all *Nostoc* MAGs. Genes required for the vanadium dependent pathway were detected in three out of six *Nostoc* species, but the pathways were potentially incomplete as the vnfH gene was not detected (Figure 2B). Syntenic analysis of orthologous regions of *Nostoc* symbiont MAGs showed large rearrangements between different genomes (Figure 2C). Cyanobacterial MAGs were shown to have significantly more transposases than other bacteria detected in the cyanolichen (Figure 2D; Table S18). More transposases were identified in Leptolyngbyaceae than Nostocaceae MAGs (Figure 2E).

### 3.3 Transposable elements drive genome expansion in chlorophyte symbiont

The chlorophyte symbiont from the tripartite cyanolichens, *Ricasolia virens* and *Lobaria pulmonaria*, was recovered independently in each sample but was shown to be the same species based on average nucleotide identity (ANI > 0.99; Table S19). Phylogenetic placement using the 18S rRNA gene showed relatedness to the trebouxioid symbiont, *Symbiochloris*. This identification was further confirmed with BLAST results (Figure 3A). The nuclear genome size was 75.8 Mb in 31 contigs (Figure 3B), with a contig N50 of 3.5 Mb, and estimated BUSCO completeness of 90.8%. Circularized organellar genomes for chloroplast and mitochondrion were recovered with sizes of 354,908bp and 105,833bp respectively (Figure S3/4).

**Figure 3:**
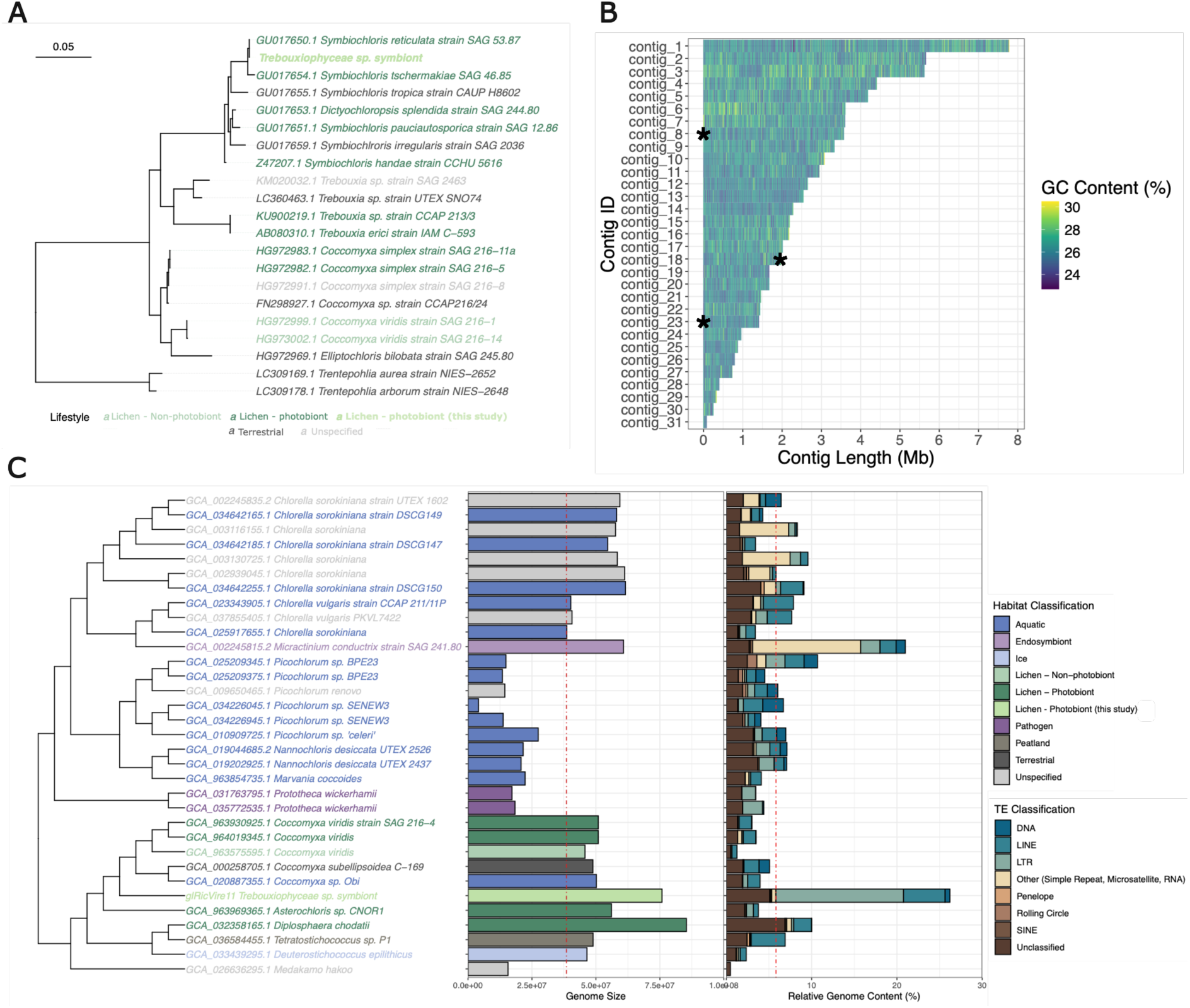
Identification and genomic characteristics of the chlorophyte photobiont in *Ricasolia virens* and *Lobaria pulmonaria*. (A) Phylogenetic tree based on the 18S rRNA locus showing taxonomic placement of recovered chlorophyte photobionts from the tripartite cyanolichens *Lobaria pulmonaria* and *Ricasolia virens* in the context of other free-living chlorophyte algae and chlorophyte symbionts. (B) GC content across contig length of the Trebouxiophyceae symbiont genome. Contig ends marked with a * indicate the detection of telomeres at the start (left-handed placement) or end (right-handed placement) of the contig. (C) Genome size and transposable element content of chlorophyte photobiont and other Trebouxiophyceae from different habitat types. Median genome size and transposable element content indicated with a dashed red line.

The genome size of the chlorophyte symbiont recovered in this study (75.8Mb) was larger than most other Trebouxiophyceae genomes (median=46.1Mb) used in our comparative analyses. Characterization of the transposable elements of the chlorophyte symbiont MAG revealed the genome to be comprised of 26% TE content (Figure 3B) and identifies this as one of the most repeat-rich Trebouxiophyceae genomes in comparison to a median TE content of 3.6% (Figure 3C). Kimura distances revealed the repeat landscape in this chlorophyte symbiont to be acquired recently in the evolutionary history of this genome, with similar Kimura distances observed in other trebouxioid symbionts and free-living strains (Figure S5).

Genes were assigned to categorical groups based on their distance to TEs ranging from overlapping (0 bp) to distant (>20,000 bp). We found 4156 genes overlapping with TE annotations in the Trebouxiophyceae symbiont from *Ricasolia virens* and *Lobaria pulmonaria* but only 65% (n = 2700) were detectably expressed (Figure 4A). Despite the same species of Trebouxiophyceae symbiont being identified in both *Ricasolia virens* and *Lobaria pulmonaria* differences in gene expression in proximity to TEs were observed between these different lichens. A lower proportion of genes, including those overlapping or in close proximity to TEs were, were observed to be detectably expressed in the Trebouxiophyceae symbiont of *Lobaria pulmonaria* (59.5%) than the symbiont of *Ricasolia virens* (77.5%) suggesting dynamic regulation of genes between individual lichen samples.

**Figure 4:**
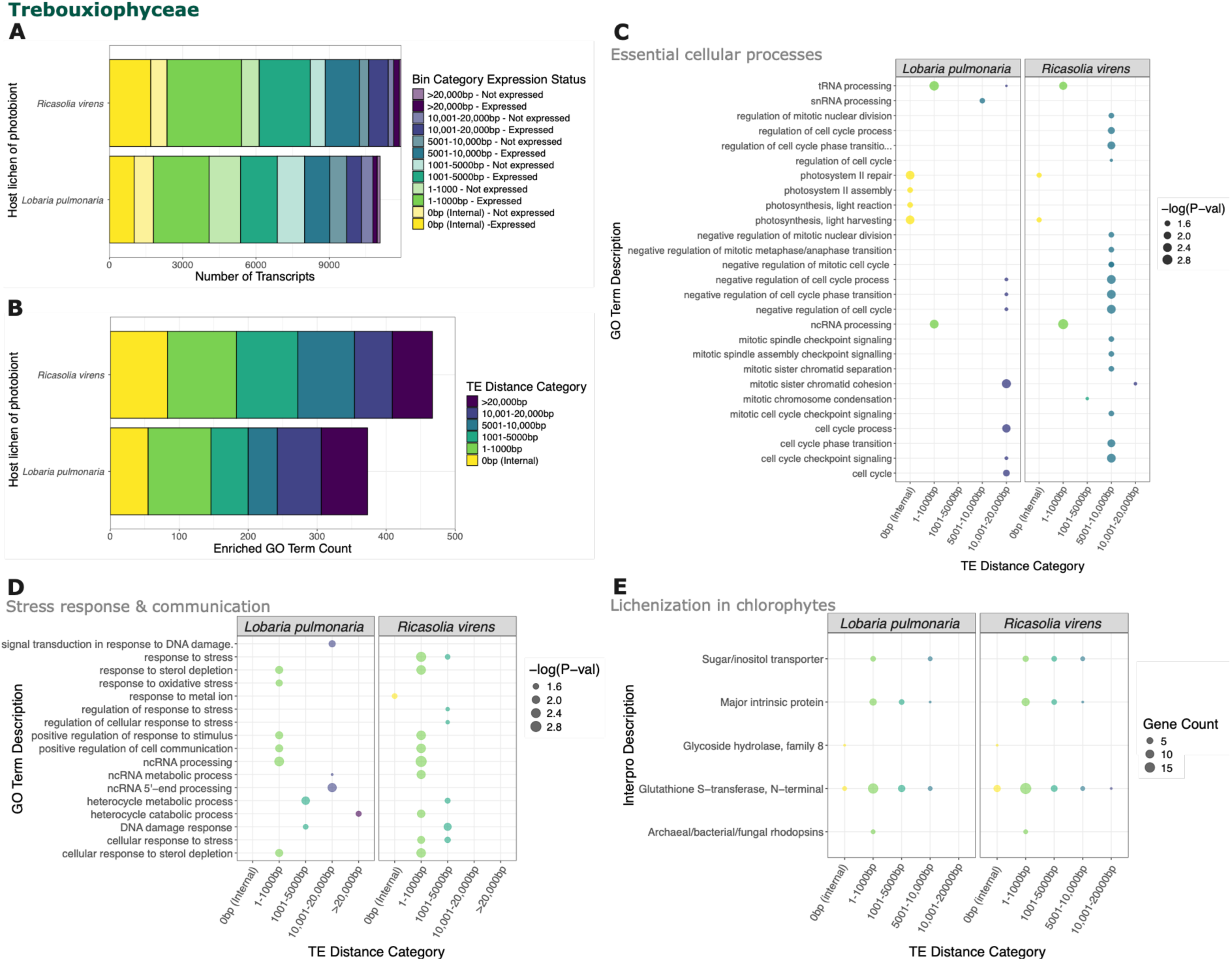
Trebouxiophyceae (*Symbiochloris*) symbiont gene expression and functional enrichment in tripartite cyanolichens, *Lobaria pulmonaria and Ricasolia virens*. (A) Expression status of genes in proximity to transposable elements shows that many genes are in close proximity to transposable elements with detectable expression in Trebouxiophyceae symbiont MAGs. (B) Frequency of enriched GO terms in Trebouxiophyceae symbiont MAGs in relation to nearest distance to transposable element. (C-D) Enrichment of GO terms in proximity to nearest TE, where size of dot represents -log(p-value). (C) Functions associated with essential cellular processes such as the cell cycle and RNA processing are observed to be enriched in genes at further distances from TE with exception of photosynthesis related GO terms. (D) Enriched GO terms show enrichment of functions corresponding to stress response and environmental communication in genes closer to transposable elements. (E) Proximity of selected genes previously associated with lichenization in chlorophytes to TEs where size of dot represents gene counts.

GO term enrichment for TE distance categories revealed variable enriched functions in Trebouxiophyceae symbionts independent of proximity to TEs and between lichens (Figure 4B; Table S20). Terms associated with essential cellular processes were found enriched at further distance from TEs with the exception of photosynthesis (Figure 4C). Stress responses and functions which may be associated with environmental communication were found to be enriched in the categorical groups corresponding to closer proximity TEs (1-1000bp; 1001-5,000bp; Figure 4D). In addition to these enriched terms, specific functions associated with chlorophyte lichenization were frequently found in close proximity to TEs (<10,000bp; Figure 4E).

### 3.4 Metagenomic recovery of largest Lecanormycetes genomes to date

Of the 15 fungal MAGs, 11 correspond to the primary fungal symbiont of the lichenized fungi from the taxonomic order Peltigerales. An additional four ascomycete fungi were identified in the cyanolichens. MAGs for the primary fungal symbionts of the Peltigerales fungi showed genomes ranging from 30.98 Mb (*Leptogium burgsessii*) to 137 Mb (*Peltigera hymenina*) (Table S9). *Peltigera* species sequenced in this study have the largest reported genome sizes for Lecanormycetes fungi characterized to date, which previously had a recorded median genome size of 66.4 Mb (Figure 5A).

**Figure 5:**
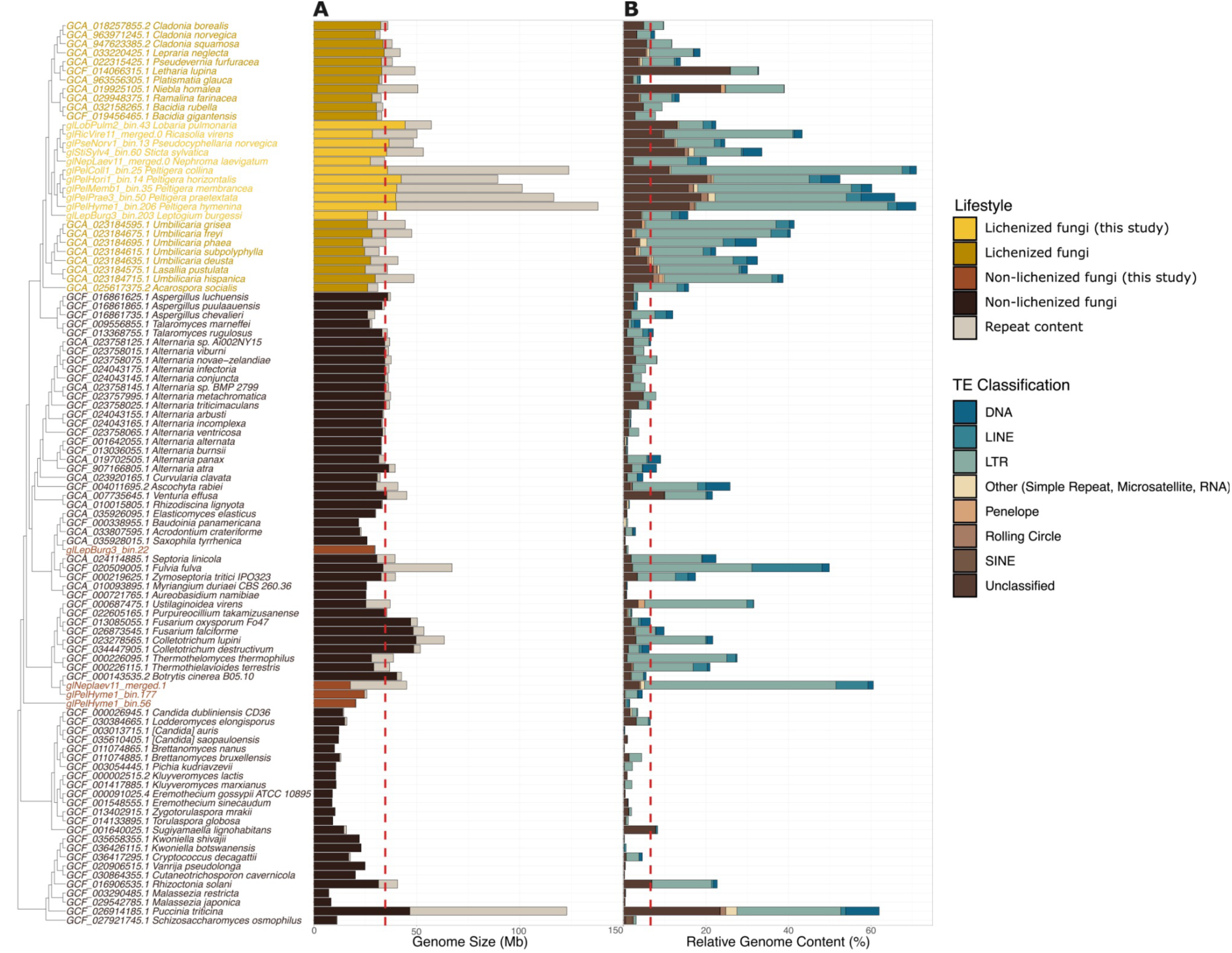
Genome size and repeat content of lichenized and non-lichenized fungi. (A) Genome size and (B) transposable element content of fungi arising from both lichenized and non-lichenized sources, including novel MAGs generated in this study. Median values of (A) genome size and (B) transposable element content (%) indicated with red dashed line.

The elevated genome sizes observed in the Peltigerales in this study may be attributed to the high TE content, which accounts for up to 71% of the genome span. These Pelitgerales are the most repeat-rich lichenized fungi genomes, and have a significantly higher repeat content than other non-lichenized fungi (Figure 5B). Contrasting the Kimura distance profiles of the TEs of the lichenized and non-lichenized fungal MAGs generated in this study showed stark differences. Peltigerales fungi have had consistent and steady TE acquisition throughout their evolutionary history, while non-lichenized fungi have comparatively shorter and smaller acquisition events (Figure S6). Transposable elements of non-lichenized fungi were generally located in broad heterochromatin-like regions, as opposed to TEs in lichenized fungi which were more commonly found in smaller TE-rich regions interspersed throughout the contigs (adjusting for genome size; Figure 6A). Gene expression levels were marginally affected by their presence within gene-rich, TE-rich, or mixed regions of the genome (Figure 6B). As a result of this pattern of TEs in the lichenized fungi genomes, many genes were found to be in close proximity to transposable elements (Figure 7A). Potential TE fusion candidates were identified based on protein domain composition (Figure S7/S8) which were shown to be actively expressed in the Peltigerales lichenized fungi (Figure S9).

**Figure 6:**
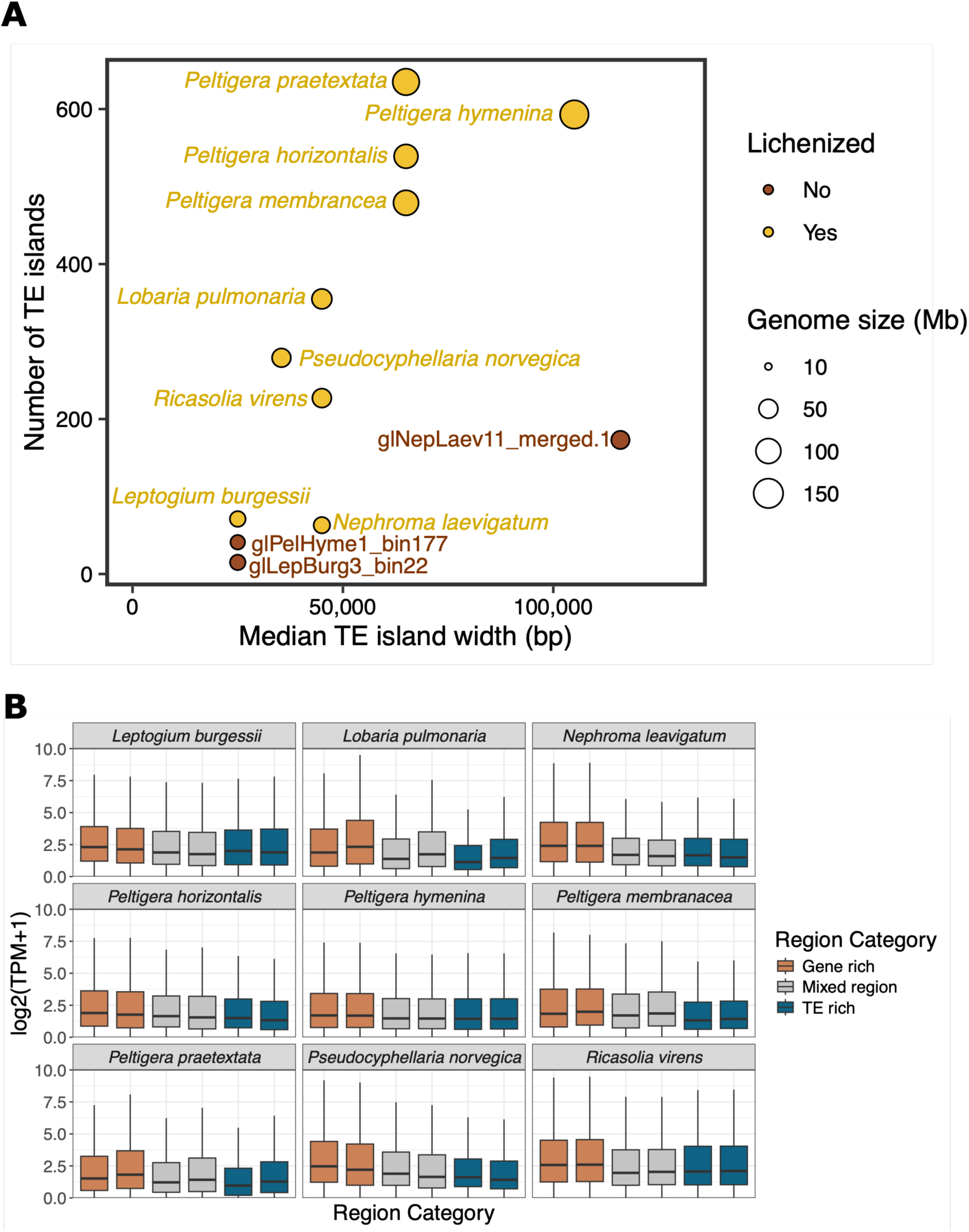
Distribution and expression of genes and TEs in fungal genomes. (A) Scatter plot showing the relationship between the number of TE-enriched regions (TE islands) and their size (TE island width), comparing with the genome size (point size) of the fungal genomes. (B) Boxplots of log TPM expression values of genes present within gene-rich, TE-rich, or mixed regions in the genomes of all sequenced lichenized fungi shown between two replicates. The average expression of genes within gene-rich regions is only slightly higher than the average expression of genes found within TE-rich regions.

**Figure 7:**
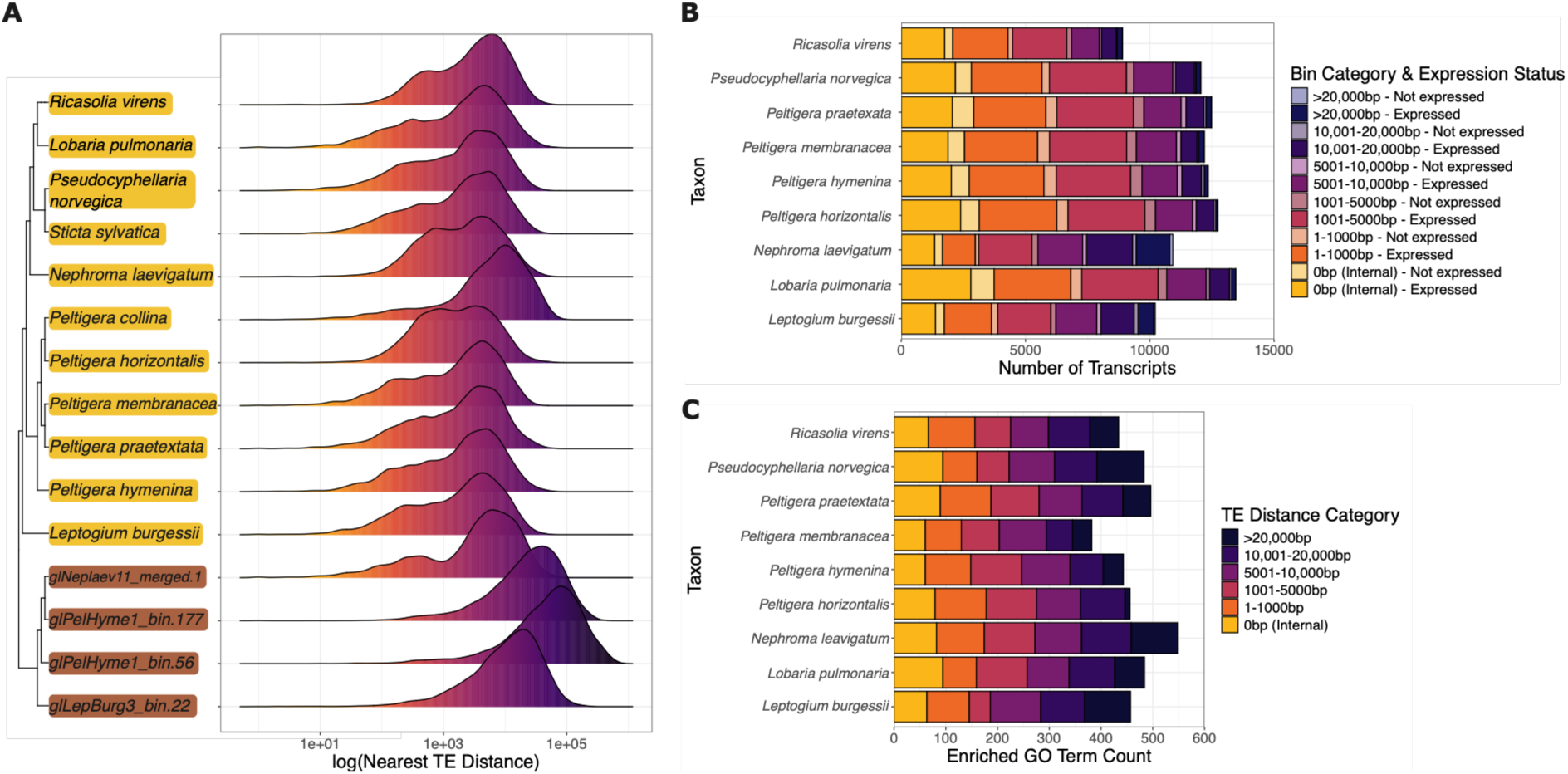
Expression and functional enrichment of genes in relation to nearest transposable element in fungal metagenomic assembled genomes (MAGs). (A) Density plot of the distance to nearest transposable element for lichenized and non-lichenized MAGs. Lichenized fungi taxa are highlighted in gold and non-lichenized in brown. (B) Expression status of genes in proximity to transposable elements shows that many genes are in close proximity to transposable elements while being actively expressed in lichenized fungal MAGs. (C) Frequency of enriched GO terms in lichenized fungi MAGs in relation to nearest distance to transposable element.

### 3.5 Cellular stress response genes enriched near transposable elements

Following annotation, 23,360 genes across 9 species of cyanolichen (Leptogium burgessii, Lobaria pulmonaria, Peltigera praetextata, Peltigera horizontalis, Peltigera hymenina, Peltigera membranacea, Ricasolia virens, Nephroma laevigatum, Pseudocyphellaria norvegica) were identified to be overlapping with transposable elements with 76% (n = 17,770) detectably expressed in RNA-seq data suggesting ongoing TE activity (Figure 7B). The closest TEs to genes that were actively expressed included DNA transposons (Fot1 and Sagan of Mariner class), long-terminal repeat (LTR) retrotransposons (Gyspy and Copia), long interspersed nuclear elements (LINE; L1 and Tad1), satellites and unclassified elements (Figure S10). DNA transposons were present significantly closer to genes on average compared to other TE families in all species, while LTR retrotransposons were significantly further away in all species (Figure S11). While a high number of genes were found to be actively expressed even when present in TE-rich islands of the genome, the overall expression of these genes was comparable, if slightly lower on average compared to genes within gene-rich regions (Figure 6B; Figure S12).

Genes were assigned to categorical groups based on their distance to TEs ranging from overlapping (0 bp) to distant (>20,000 bp). GO term enrichment for TE distance categories revealed enriched functions in fungal genomes dependent on proximity to TE (Figure 7C; Table S21). Genes, globally, were further away from TEs than expected by chance (Table S22). Despite this, genes overlapping or in close proximity (<1000 bp) to TEs were found to be enriched for cellular responses to stress and stimulus (e.g., radiation, UV, abiotic stress), transmembrane transport, and biosynthesis and metabolism of secondary metabolites including mycotoxins and heterocycles (Figure 8A). Genes at a greater distance from TEs were enriched for GO terms associated with essential cellular processes such as RNA processing and cell cycle maintenance (Figure 8B), suggesting stronger selection against TE invasion of such genes. Similar results were obtained using permutation tests, shuffling the locations of the genes to determine whether any GO terms were associated with genes closer or further away from TEs than expected (Figure S13). Semantic enrichment and clustering revealed some distinct terms enriched between those significantly closer or further away from TEs (Figure S14/S15).

**Figure 8:**
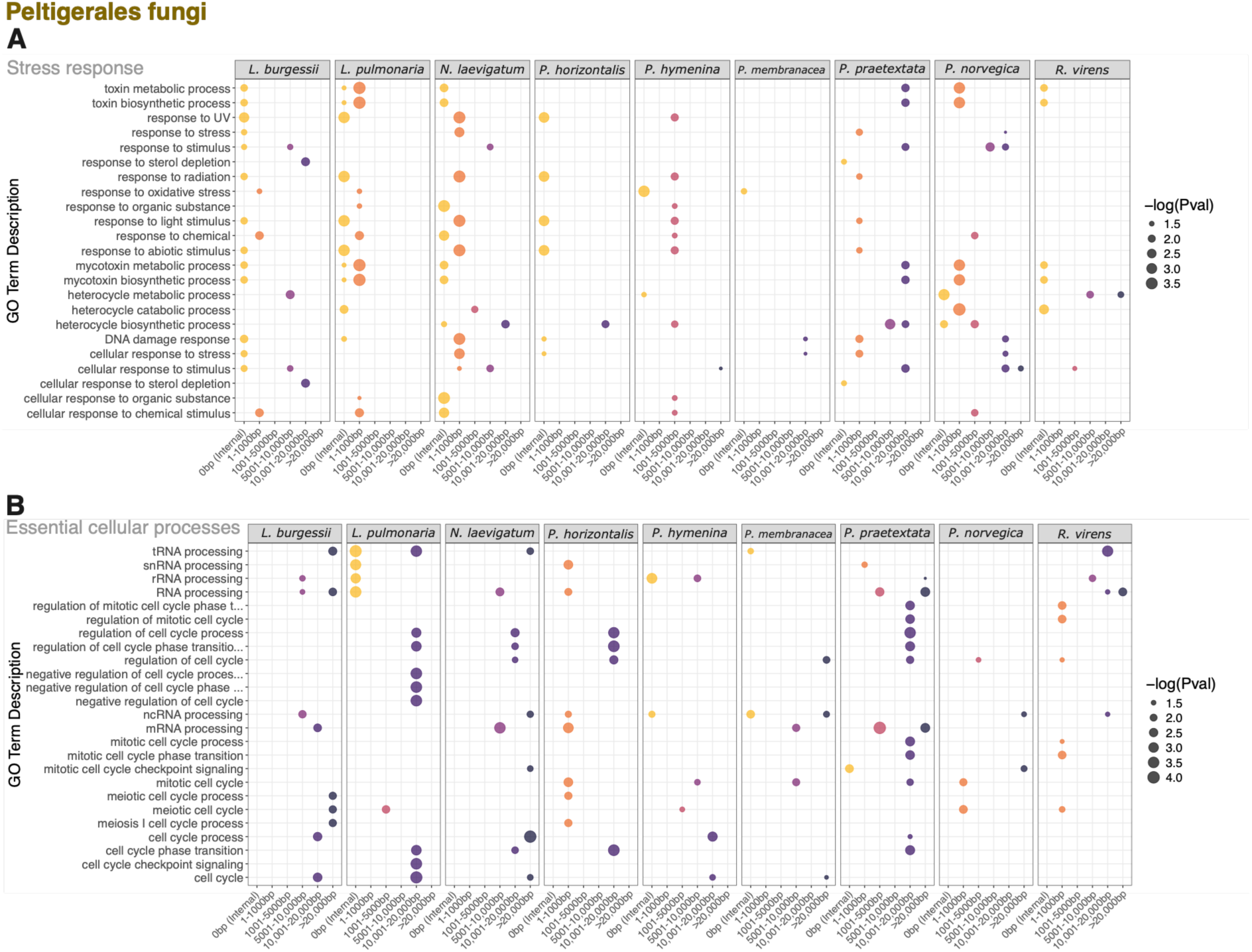
Enriched GO terms in Peltigerales fungal metagenomic assembled genomes (MAGs). Enriched GO terms corresponding to stress response and environmental communication (A) and essential cellular processes (B) where the point size represents -log(p-value). GO terms corresponding to stress response were observed in genes closer to TEs while those associated with essential cellular processes were at further distances from TEs.

## 4. Discussion

We performed long-read metagenomic sequencing of 11 species of cyanolichens to explore organismal composition and genomic content in the symbiotic association. The lichen-associated bacterial communities were found to contain key lineages that have previously been reported in lichens^9^ (Figure 1G). We found frequent co-occurrence of multiple species of *Nostoc* within the same lichen (Figure 1I) emphasising previous demonstrations of cyanobacterial photobiont distribution and promiscuity^7,66,67^. Some *Nostoc* symbionts were also likely to possess an alternative vanadium dependent nitrogen fixation pathway (Figure 2B) as previously observed in other cyanobacterial symbionts^7,68–70^ suggesting a broader specialization to support nitrogen fixation independent of environmental conditions. Genomic rearrangements and high counts of transposase genes in the cyanobacterial genomes (Figure 2D) also suggest the potential impact of mobile genetic elements in essential bacterial symbionts to allow for host adaptation^71^.

The ability to explore the genomics of the eukaryotic symbionts in lichens has frequently lagged behind the exploration of their bacterial counterparts due to the challenge of recovering high-quality, highly contiguous MAGs from short-read metagenomic datasets. Despite the bulk of biomass arising from the fungal partner, previous metagenomic studies using short-read sequencing have faced challenges in recovery of both fungal and algal MAGs^9^. This has prevented advances in the understanding of lichen evolution as the number of publicly available lichenized fungal genomes differs by an order of magnitude from current estimated species counts. In this study, we generated highly contiguous, high quality fungal genomes for 15 fungi, 11 of which are lichenized, presenting the opportunity to explore genomic characteristics and specialized molecular functions in the lichenized lineage, Peltigerales.

The dependence of lichenized fungi on their photobiont for growth draws parallels to the lifestyle features observed in obligate biotrophic pathogenic fungi, which can only grow on living host tissue. We observe similar trends in genome size and TE content between these two groups of fungi with co-dependent lifestyles^20^. Many of the largest fungal genomes characterized to date arise from obligate biotrophic pathogens with the largest, *Austropuccinia psidii* with a genome of 1 Gb, being composed of >90% TEs^19^. While not as extreme, the Peltigerales lichens sequenced in this study have above-average genome sizes that can be attributed to high TE content and represent the most TE-rich genomes of lichenized fungi characterised to date (Figure 5). Transposable element content in the context of fungal lifestyles has been explored previously^72,73^ and the trends observed here add a further layer to the impact of lifestyle on TE content: the degree of dependency of fungi on other organisms (e.g. obligate lifestyles) appears to be associated with larger genome sizes driven by increased TE content.

The biological impact of increased numbers of TEs and their distribution in a genome is likely complex. TEs can influence the expression of genes near insertion sites, by adding potential promoters and enhancers, and by impacting expression in local domains when they are epigenetically silenced. We explored whether genes with annotations overlapping or near TE annotations were more or less likely to be expressed. We confirm that genes overlapping with and nearby to TEs in lichenized fungi are actively expressed at levels comparable to other genes and that these genes can be associated with functions with biological relevance to the lichenized lifestyle (Figure 8A). These results also evoke a similar phenomenon observed in fungal pathogen TE evolution, where the copy number and diversity of actively expressed virulence genes (often present near TEs) is positively associated with TE expansion occurring in smaller TE-rich regions scattered throughout the genome^20^. This parallel could suggest that the ability of Peltigerales lichens to participate in the symbiosis may be linked to such TE proximal genes undergoing rapid evolution. When looking into genes in close proximity to TEs in the Peltigerales lichens, we observed a high abundance of genes associated with responses to stress, environmental stimuli and production of secondary metabolites which correspond to many of the characteristics associated with lichenization and their ability to survive harsh environments and produce diverse secondary metabolites. The samples used in this study were desiccated and the transcriptomes reflected this metabolic status. Future transcriptomic studies exploring the life cycle and desiccation processes in lichens will be required to further elucidate the impact of TEs on gene regulation.

In the gradient of necessity of the symbiotic relationship in lichens, the lichenized fungi within the Lecanoromycetes lean towards being obligate symbionts, leaving the question remaining as to whether similar patterns of TE content and genome expansion are observed in non-obligate symbionts, such as lichenized chlorophytes or other fungal lineages. The TE content of algae has been underexplored with estimates of TEs contributing to a small portion of the overall genome size in the Chlorophyta. In our analysis of the Trebouxiophyceae symbiont, detected in both *Ricasolia virens* and *Lobaria pulmonaria*, we show that some lichenized trebouxioid photobionts may have elevated TE content, but this is often comparable to that observed in free-living aquatic species. Recent research on lichenization in chlorophytes revealed specialized carbohydrate active enzymes amongst other functions associated with adaptation to high light intensity and drought tolerance^8,9^. In our analyses, genes associated with these functions were frequently found in close proximity to TEs (Figure 4E; Figure S15). A carbohydrate active enzyme from glycoside hydrolase family 8 that has been associated with the lichenization of chlorophytes^8^ was noted to contain a TE fragment within it. While this may play a role in the functional adaptation of the algae, the absence of significant genome expansions as observed in the lichenized fungi presented in this work could indicate one of two scenarios: either the Trebouxiophyceae symbiont is under strong pressure to control TE content and does not accept TE insertions, or the Trebouxiophyceae symbiont is under less pressure to adapt and fewer genomic changes, including TE expansions, are occurring.

The relationship of genes and TEs in lichen symbionts can broadly be classified into five simplified scenarios: (1) genes are arising from domesticated TEs; (2) TEs are impacting the expression of functional genes relevant to symbiosis and the lichenized lifestyle; (3) novel functional genes essential to the lichenized lifestyle are formed through the fusion of TE domains to genome domains; (4) genes are not impacted by TE activity; or (5) the distribution of smaller TE-rich regions in the genome is leading to an increase in copy number variation of symbiosis-related genes. The work presented here constitutes the first exploration of TE expression and activity in the Peltigerales fungi and their associated chlorophyte photobionts and identifies a potential association of obligate versus facultative lifestyle on high versus low TE content and TE distribution. As more metagenomic research utilises long-read sequencing, the expected recovery of high-quality, highly-contiguous eukaryotic MAGs will increase creating further opportunity to characterize TE content in diverse microbial eukaryotes and population level genome dynamics to further advance our understanding of these genomic elements on evolutionary processes.

## Acknowledgements

We acknowledge the support of NatureScot for site access and sampling permissions for field samples collected in Glen Creran Woods. This work was supported by EMBL (European Molecular Biology Laboratory) core funds and Wellcome Sanger Institute (220540/Z/20/A). This research was funded in part, by the Wellcome Trust. For the purpose of Open Access, the author has applied a CC BY public copyright licence to any Author Accepted Manuscript version arising from this submission.

## Data and code availability

The raw sequencing data generated in this project at the Wellcome Sanger Institute is publicly available under project accessions PRJEB63593 and PRJEB73802. Reference sequences used for phylogenetic placement and comparison available in public repositories with accession information in Supplementary Tables S12, S14, S15, S16. Custom scripts for data analysis and visualization are available at https://github.com/Finn-Lab/2026_transposable_elements_cyanolichens. Additional files required to reanalyze the data including annotation files and associated data frames available upon request.

