## Supplementary material for "Transposable Element Diversification and the Evolution of Peltigerales Lichen Symbionts": Figure S1

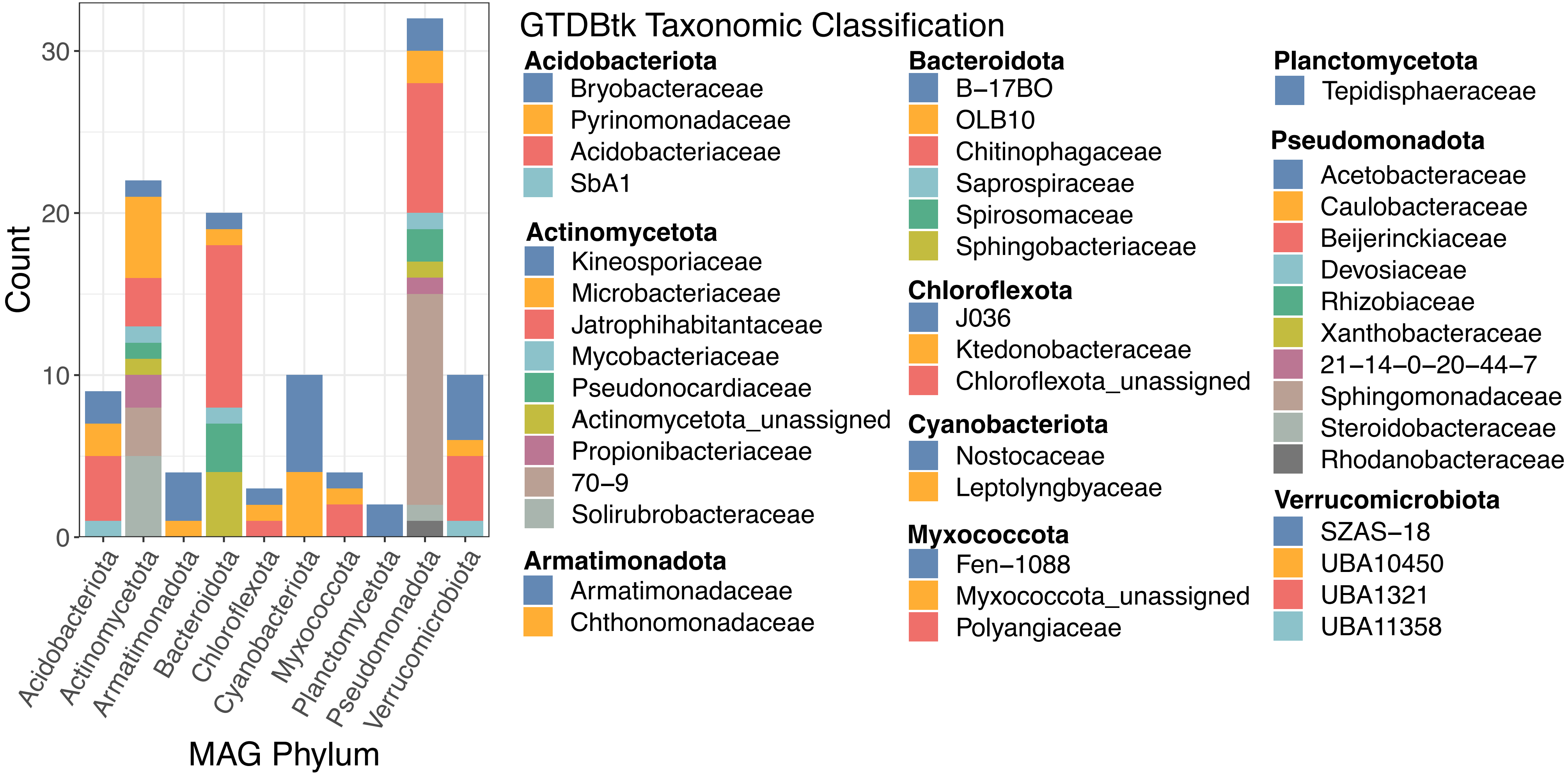


**Figure S1:** GTDBtk taxonomic family classification of dereplicated bacterial metagenomic assembled genomes (MAGs) detected in cyanolichens. A large number of MAGs were found to belong to the phyla Pseudomonadota, Actinomycetota and Bacteroidota.
