## Supplementary material for "Transposable Element Diversification and the Evolution of Peltigerales Lichen Symbionts": Figure S2

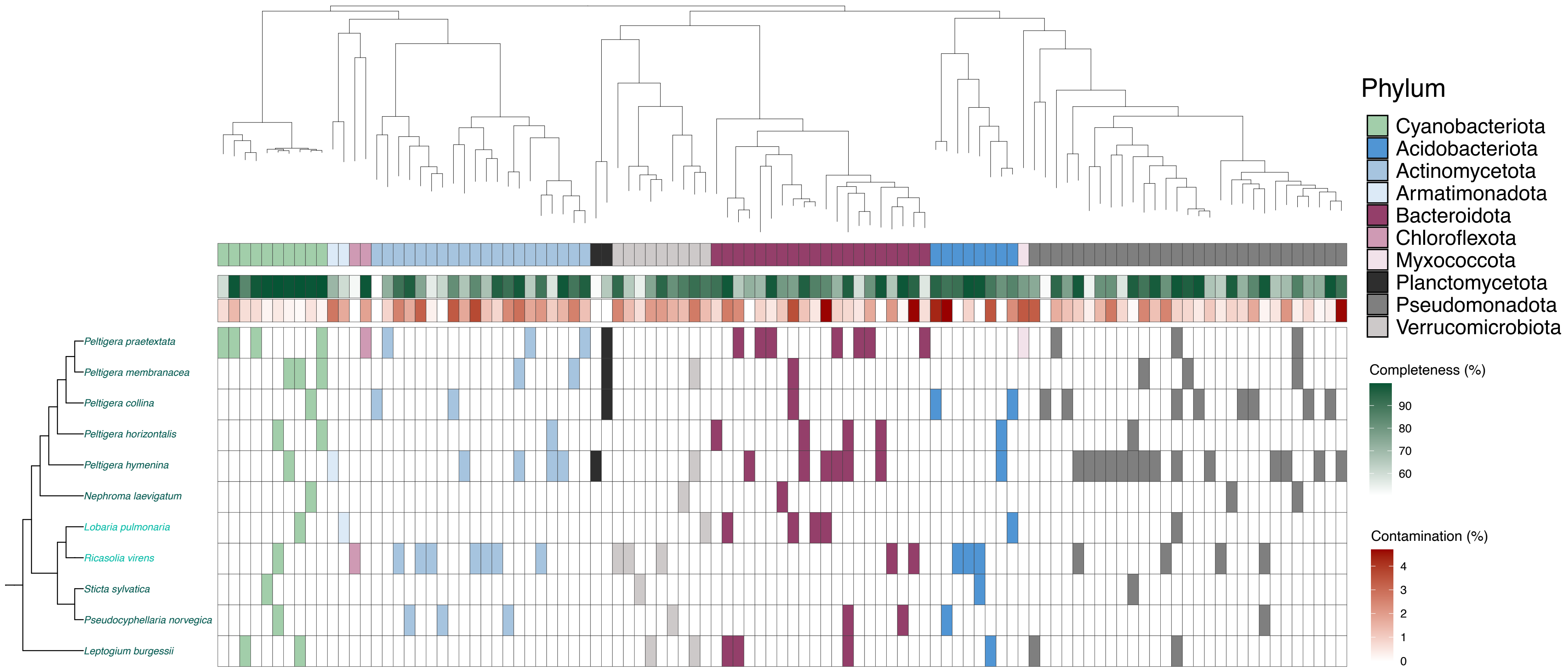


**Figure S2:** Occurrence of dereplicated bacterial metagenomic assembled genomes (MAGs) across cyanolichens. Presence of MAG in a cyanolichen indicated by shaded tiles coloured according to phylum. Completeness and contamination of MAGs as calculated with CheckM shown with gradient shading. Tripartite cyanolichens are indicated with text labels in bright turquoise.
