## Supplementary material for "Transposable Element Diversification and the Evolution of Peltigerales Lichen Symbionts": Figure S3

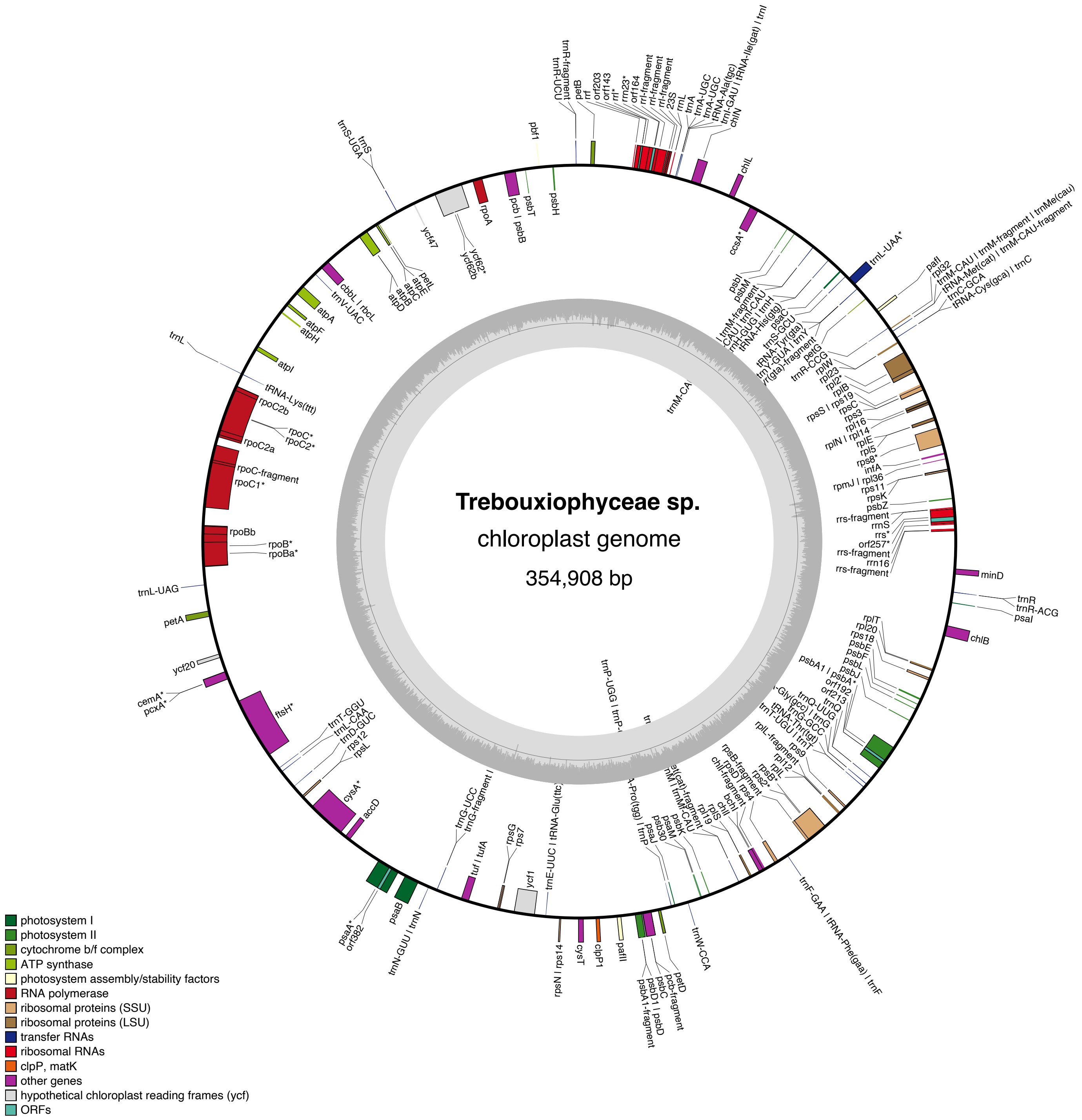


**Figure S3:** Chloroplast genome map for Trebouxiophyceae photobiont detected in tripartite cyanolichens, *Lobaria pulmonaria* and *Ricasolia virens*.
