## Supplementary material for "Transposable Element Diversification and the Evolution of Peltigerales Lichen Symbionts": Figure S4

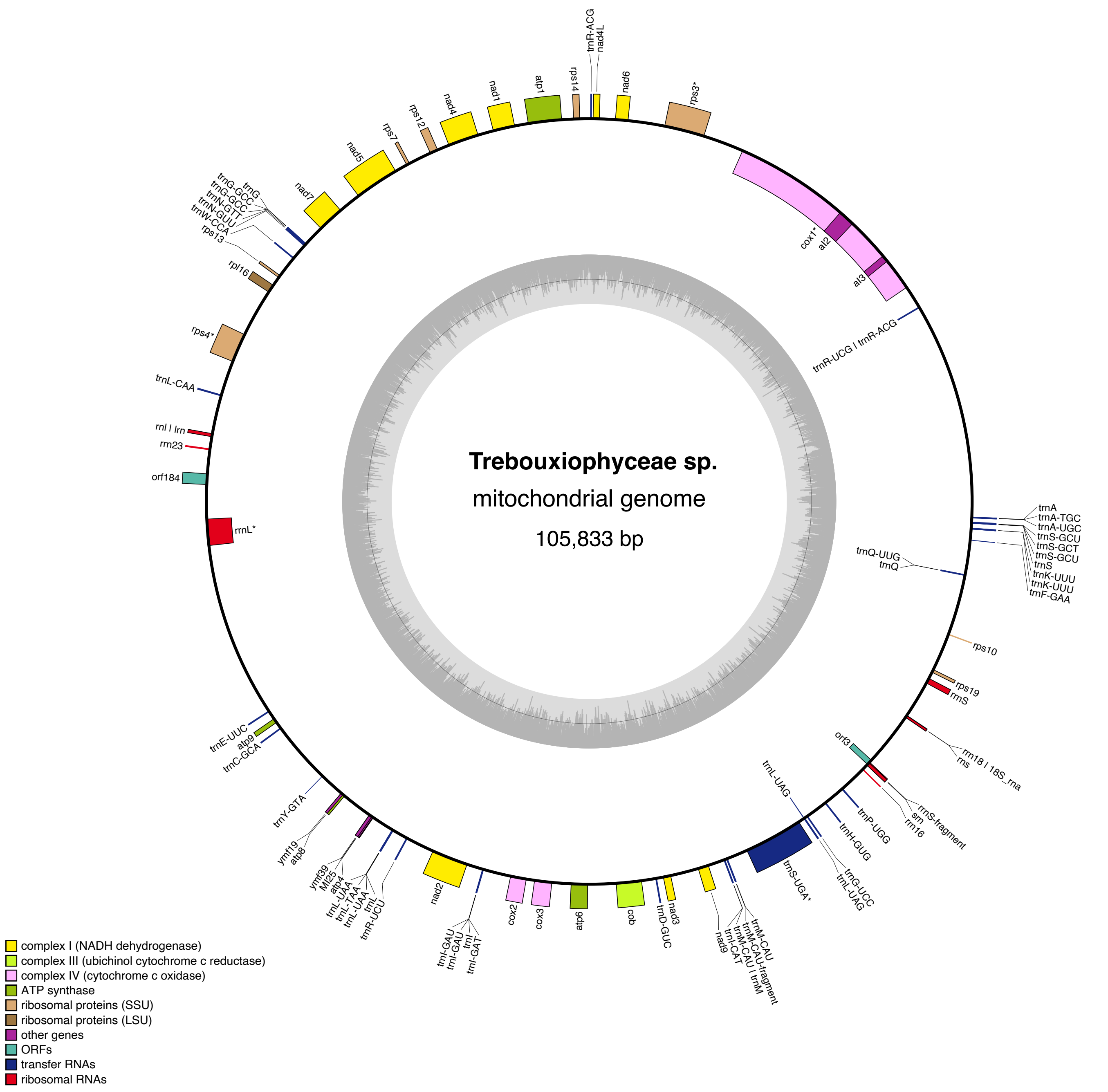


**Figure S4:** Mitochondrion genome map for Trebouxiophyceae photobiont detected in tripartite cyanolichens, *Lobaria pulmonaria* and *Ricasolia virens*.
