## Supplementary material for "Transposable Element Diversification and the Evolution of Peltigerales Lichen Symbionts": Figure S5

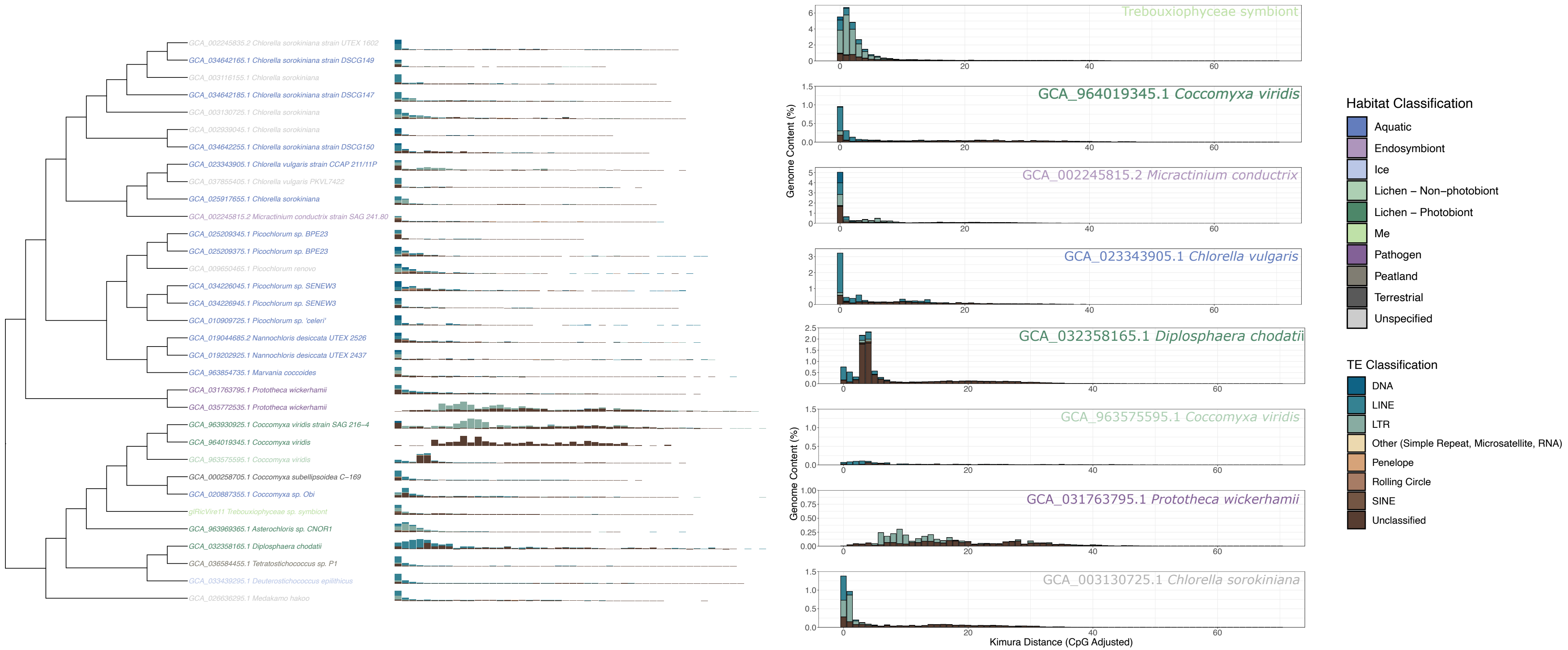


**Figure S5:** Transposable element landscape of trebouxoid chlorophyte algae from diverse habitats including the Trebouxiophyceae symbiont generated in this study, other lichens and free-living sources. Kimura distances show TE acquisition events varied across habitat types with some experiencing recent acquisition events in their evolutionary history.
