## Supplementary material for "Transposable Element Diversification and the Evolution of Peltigerales Lichen Symbionts": Figure S6

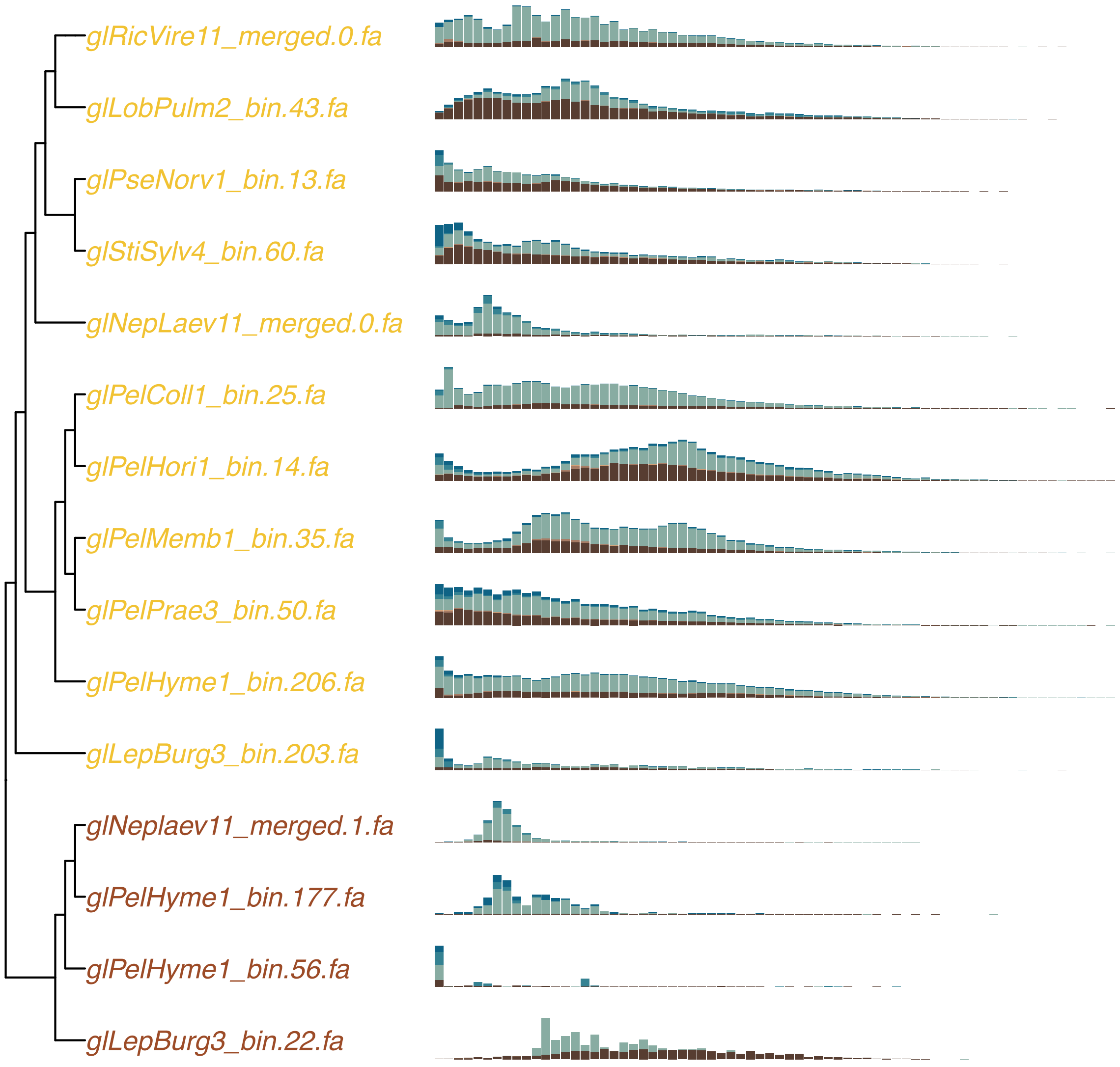


**Figure S6:** Transposable element landscape of fungal metagenomic assembled genomes (MAGs) generated in this study. Lichenized fungal MAGs, indicated with yellow text labels, showed similar TE acquisition events based on Kimura distances.
