## Supplementary material for "Transposable Element Diversification and the Evolution of Peltigerales Lichen Symbionts": Figure S7

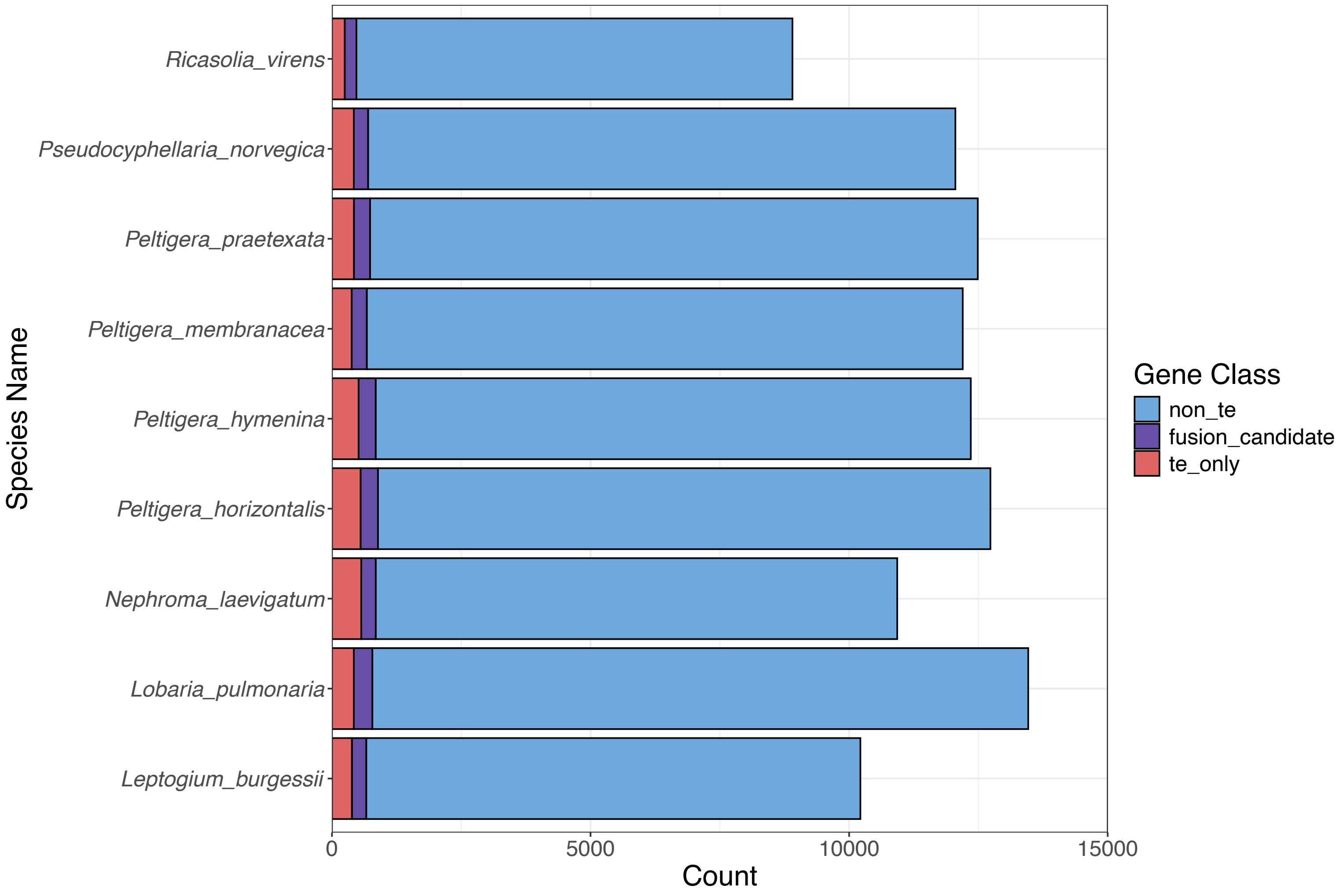


**Figure S7:** Count of genes predicted to be from transposable elements (TE) and TE fusion candidates in lichenized fungal metagenomic assembled genomes generated in this study.
