## Supplementary material for "Transposable Element Diversification and the Evolution of Peltigerales Lichen Symbionts": Figure S8

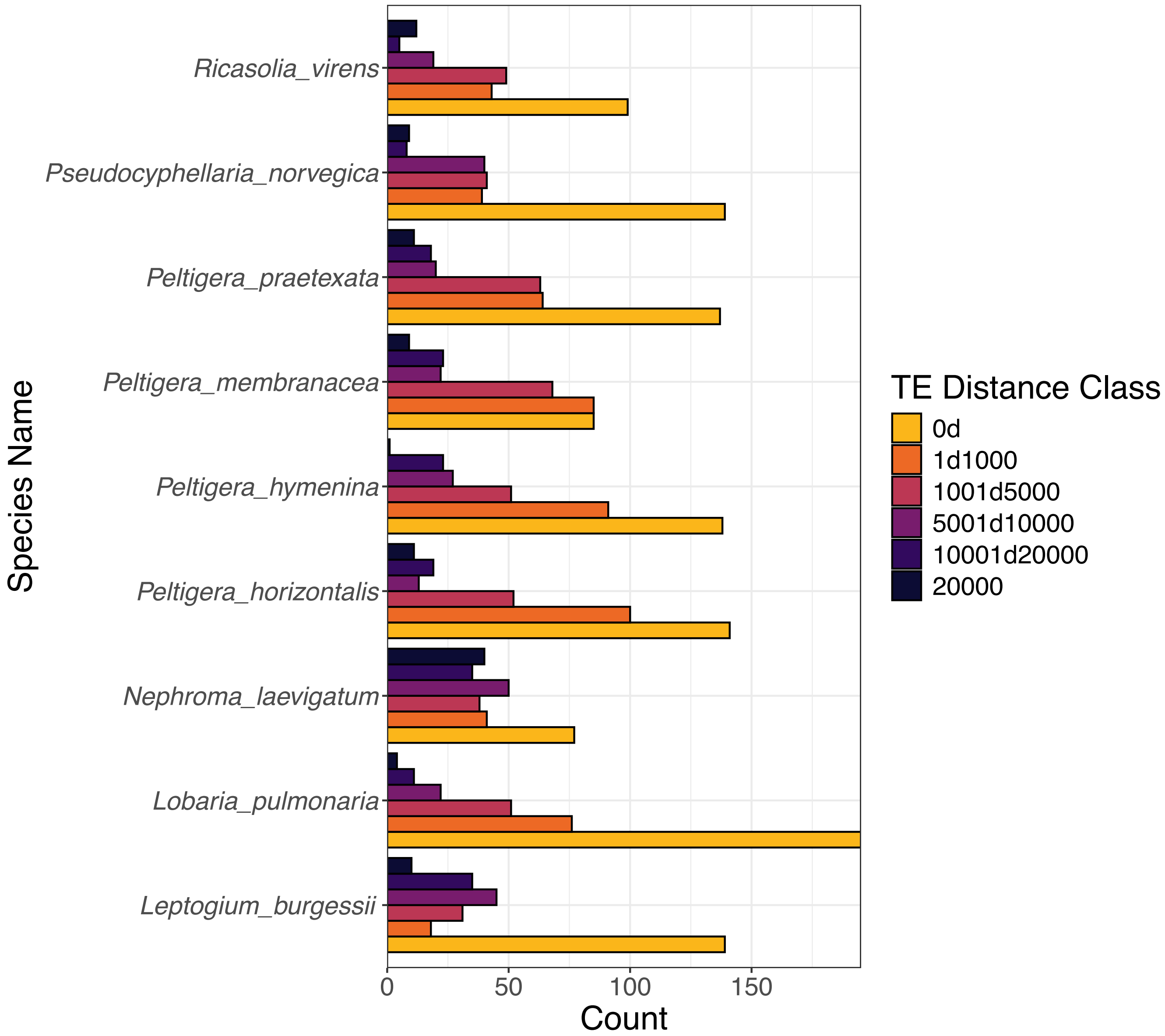


**Figure S8:** Distance of transposable element (TE) fusion gene candidates in lichenized fungal metagenomic assembled genomes to TE reveals a higher proportion of fusion candidates in closer proximity to TEs.
