## Supplementary material for "Transposable Element Diversification and the Evolution of Peltigerales Lichen Symbionts": Figure S9

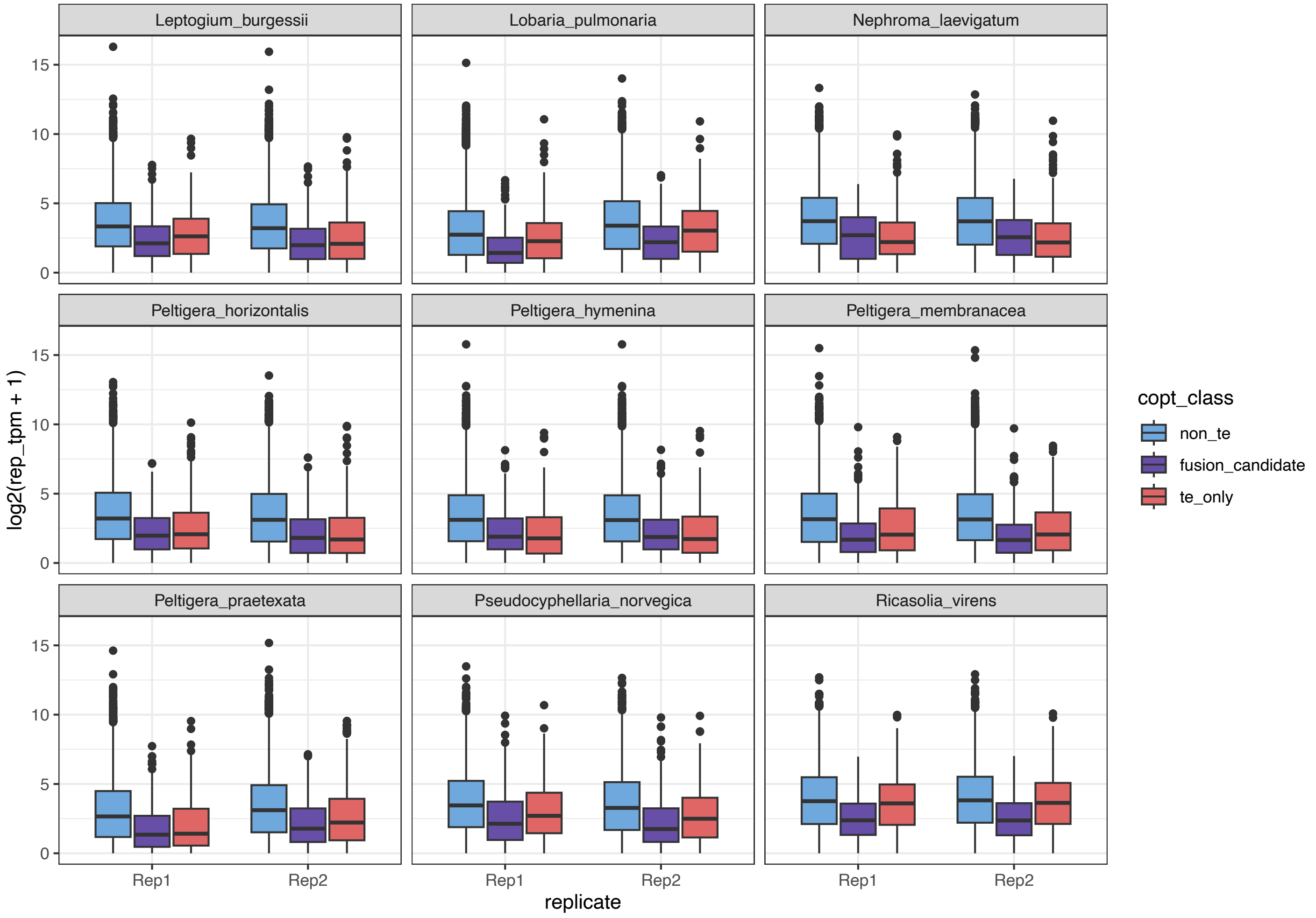


**Figure S9:** Expression levels of transposable elements (TE), non-TE and TE fusion candidate genes in lichenized fungal metagenomic assembled genomes assessed in this study.
