## Supplementary material for "Transposable Element Diversification and the Evolution of Peltigerales Lichen Symbionts": Figure S10

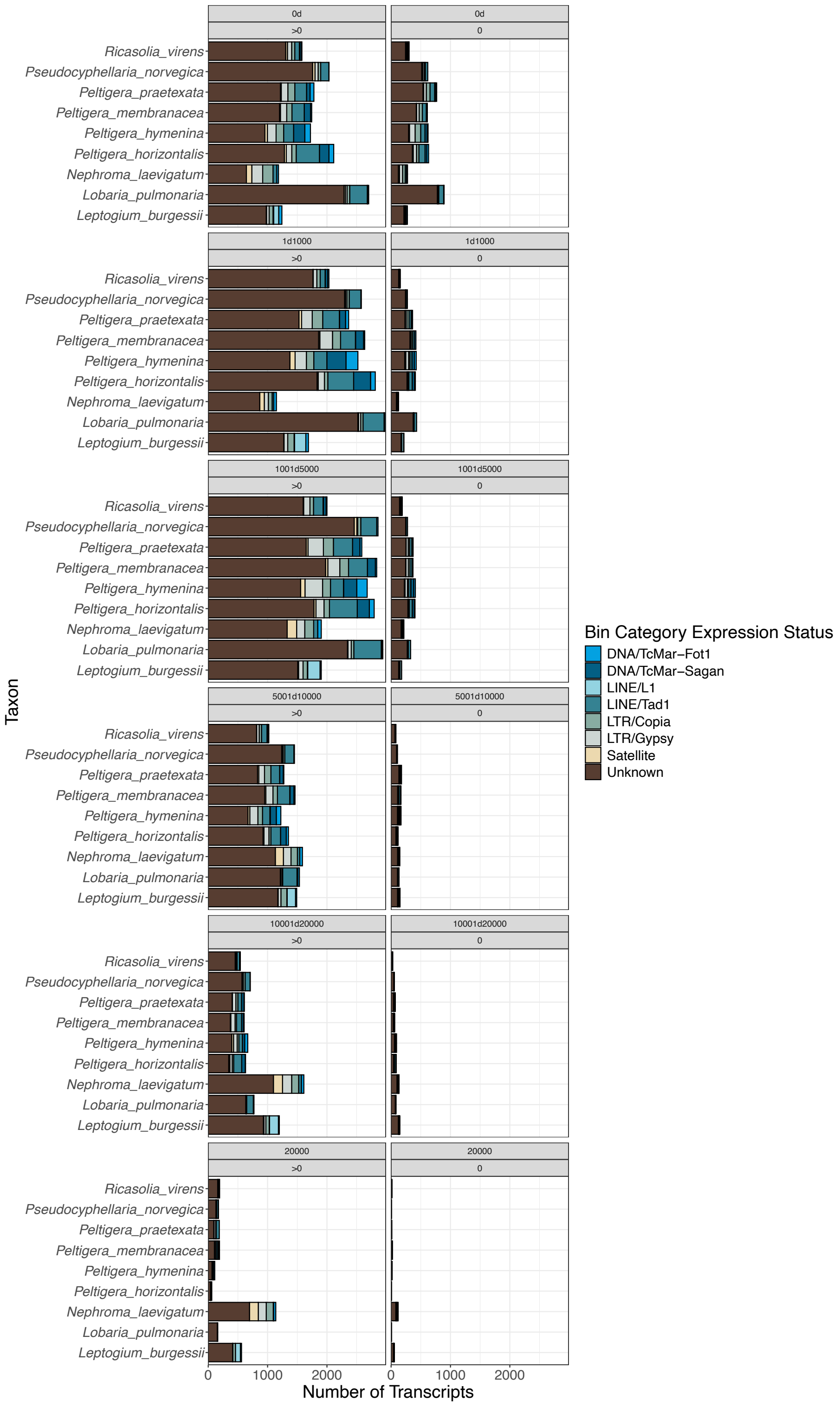


**Figure S10:** Count of genes in lichenized fungal metagenomic assembled genomes expressed (>0) and not expressed (0) in relation to proximity to transposable elements (TE) and class of nearest TE.
