## Supplementary material for "Transposable Element Diversification and the Evolution of Peltigerales Lichen Symbionts": Figure S11

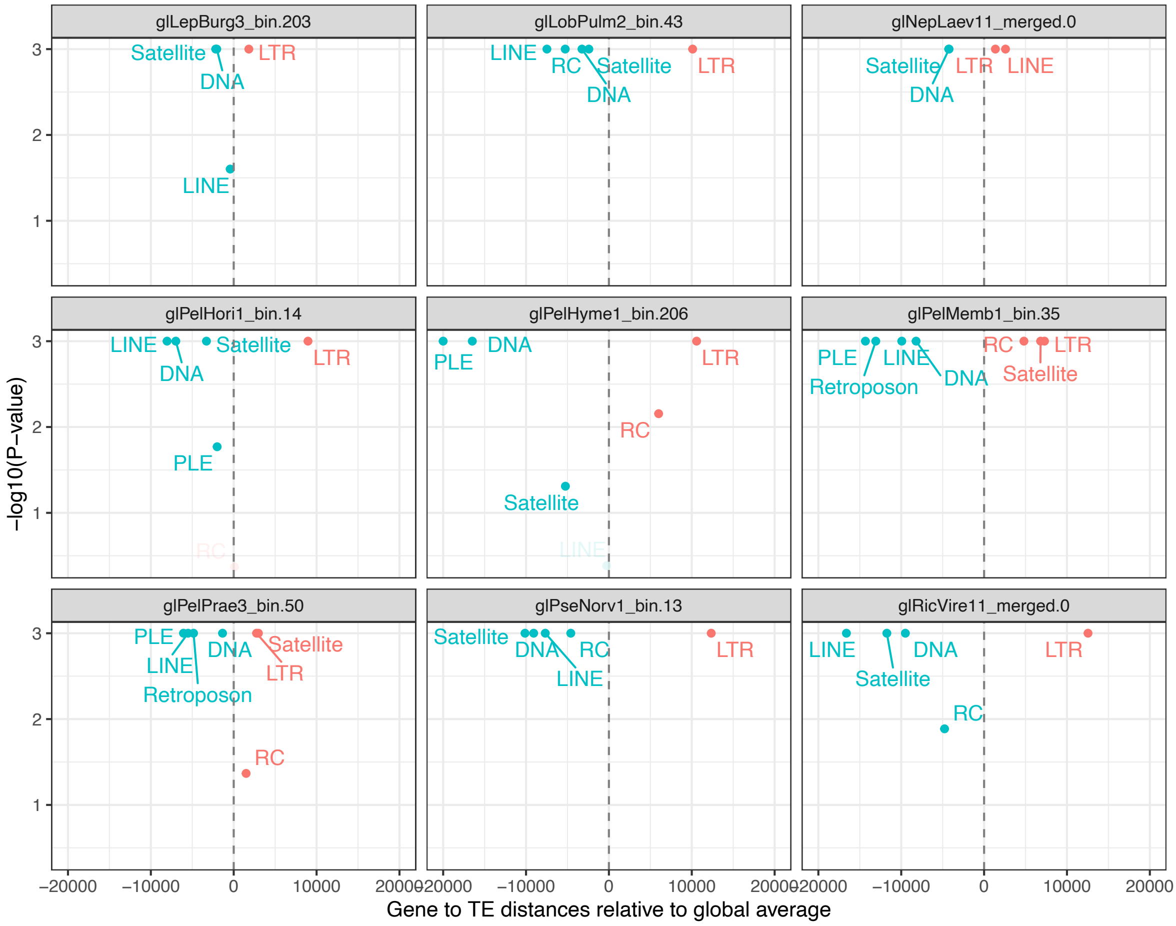


**Figure S11:** Specific families of transposable elements (TE) show non-random distributions in the genomes of lichenized fungi. Using permutation testing, TE families were shuffled 1000 times and P-values calculated based on the observed distance between specific families of TEs and genes. The x-axis shows the average distance of each TE family relative to the global average distance of all TEs with an annotated family, while the y-axis shows the negative log10-transformed P-value. Non-significant points ($P > 0.05$) are plotted with a lighter colour.
