## Supplementary material for "Transposable Element Diversification and the Evolution of Peltigerales Lichen Symbionts": Figure S12

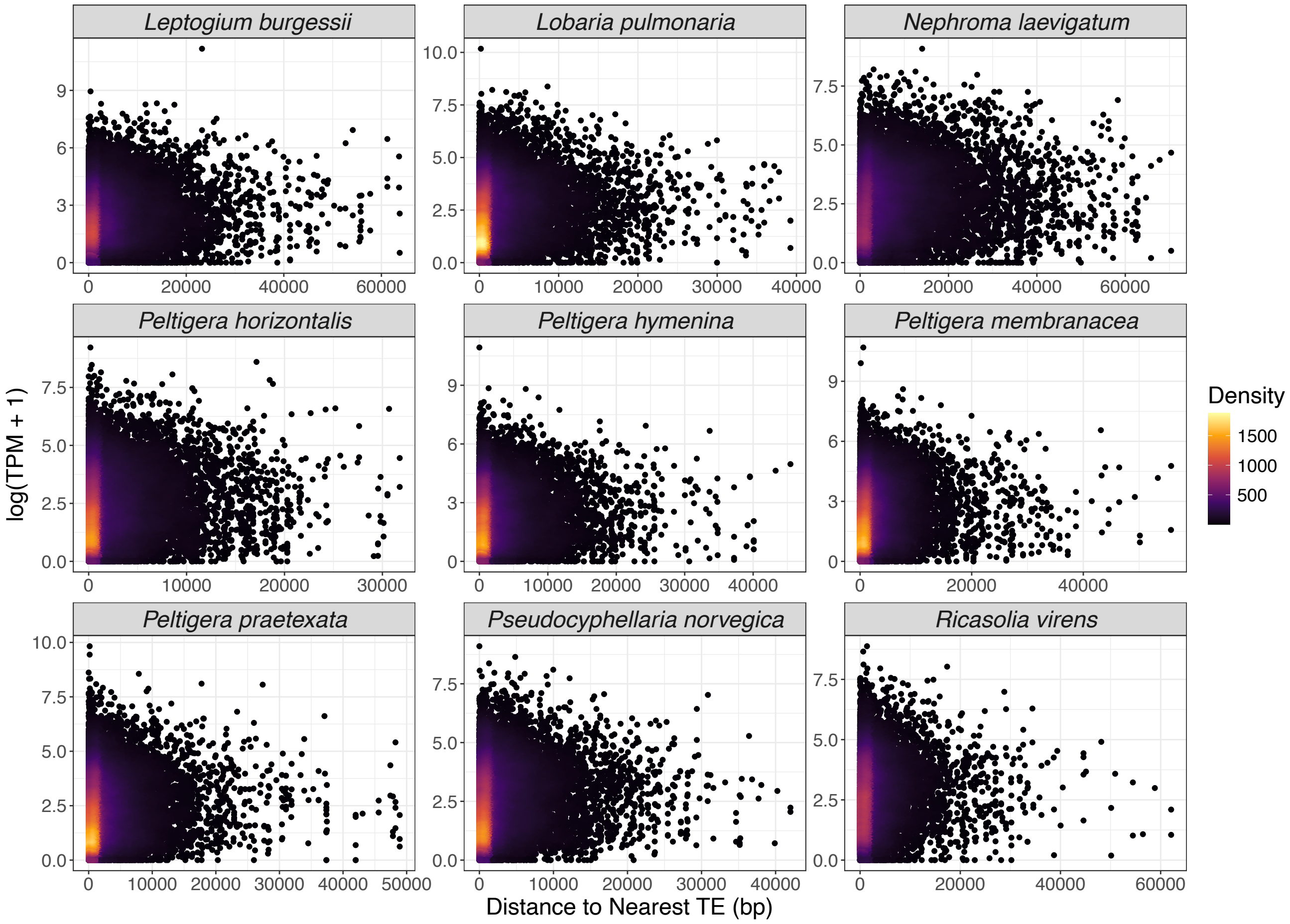


**Figure S12:** Gene expression levels are not strongly affected by their proximity to TEs. For each of the lichenized fungi, genes were plotted by their distance to the nearest TE and their log10-transformed expression (TPM).
