## Supplementary material for "Transposable Element Diversification and the Evolution of Peltigerales Lichen Symbionts": Figure S13

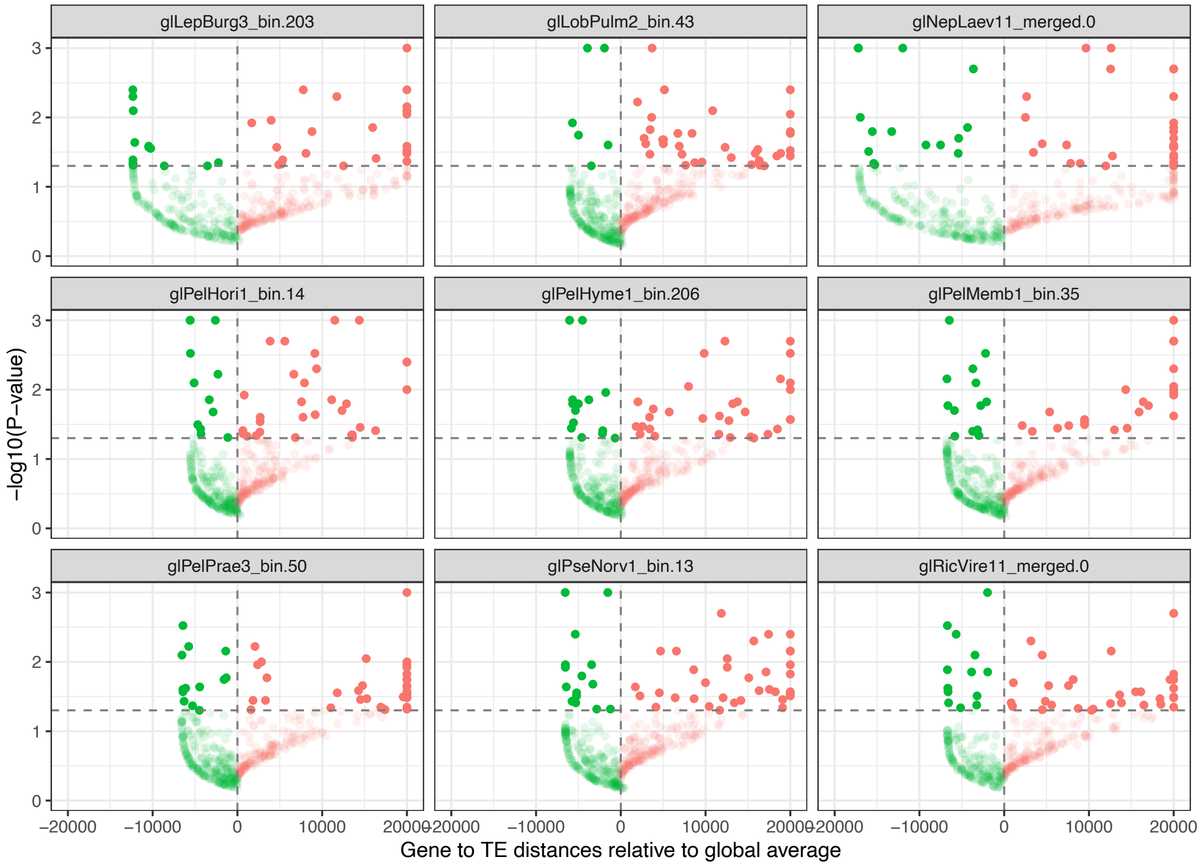


**Figure S13:** Certain Gene Ontology (GO) Biological Process (BP) terms are non-randomly distributed in the genomes of lichenized fungi relative to their distance to TEs. Using permutation testing, GO:BP terms of annotated genes were shuffled 1000 times and P-values calculated based on the observed distance between genes associated with specific terms and TEs. The x-axis shows the average distance of genes grouped by GO:BP term relative to the global average distance of all annotated genes, while the y-axis shows the negative log10-transformed P-value. Non-significant points ($P > 0.05$) are plotted with a lighter colour.
