## Supplementary material for "Transposable Element Diversification and the Evolution of Peltigerales Lichen Symbionts": Figure S14

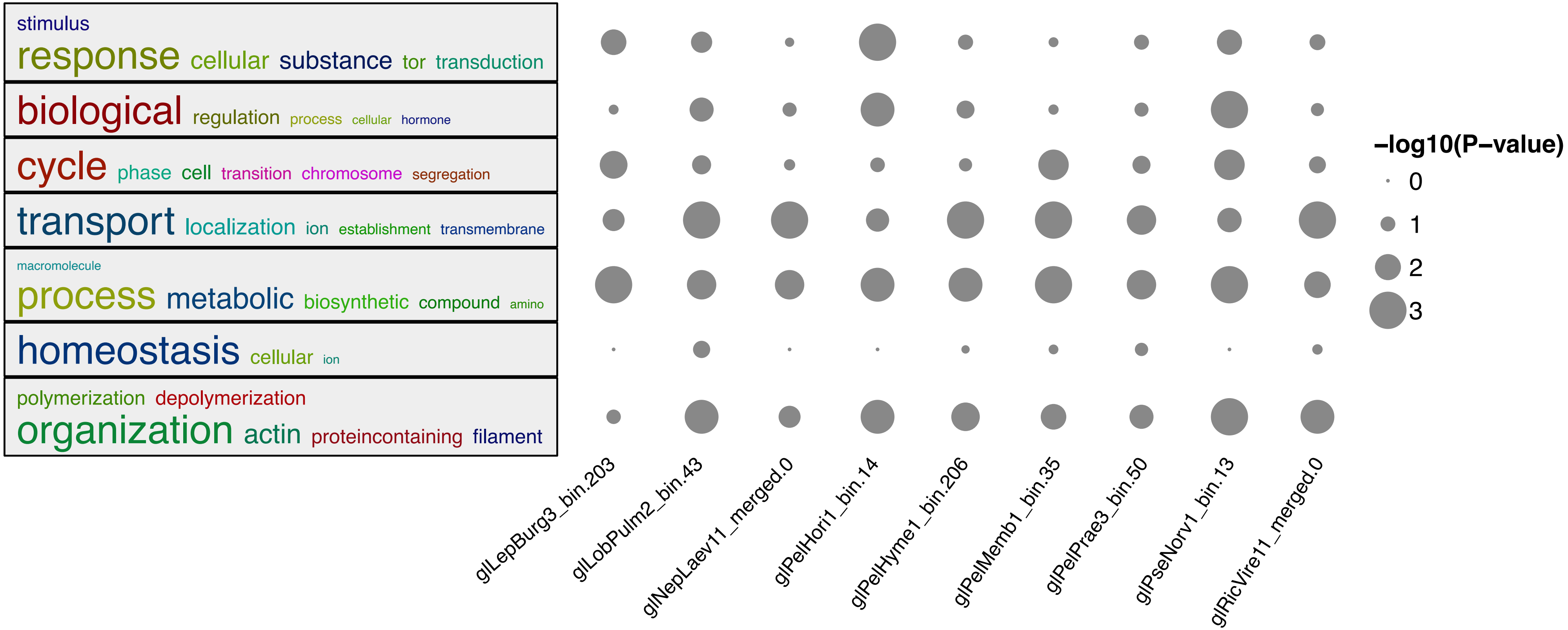


**Figure S14:** Semantic enrichment and clustering of GO:BP terms with an average gene distance further away than the global average distance (Figure S13). The most significant term was plotted for each sample and cluster using their negative log10-transformed P-value.
